# Unbiased identification of cell identity in dense mixed neural cultures

**DOI:** 10.1101/2024.01.06.574474

**Authors:** Sarah De Beuckeleer, Tim Van De Looverbosch, Johanna Van Den Daele, Peter Ponsaerts, Winnok H. De Vos

**Affiliations:** Laboratory of Cell Biology and Histology, University of Antwerp; Laboratory of Experimental Hematology, Vaccine and Infectious Disease Institute (Vaxinfectio), University of Antwerp; Antwerp Centre for Advanced Microscopy, University of Antwerp; µNeuro Research Centre of Excellence, University of Antwerp

**Keywords:** morphological phenotyping, multiplexed imaging, iPSC culture validation, neural differentiation, computational biology

## Abstract

Induced pluripotent stem cell (iPSC) technology is revolutionizing cell biology. However, the variability between individual iPSC lines and the lack of efficient technology to comprehensively characterize iPSC-derived cell types hinder its adoption in routine preclinical screening settings. To facilitate the validation of iPSC-derived cell culture composition, we have implemented an imaging assay based on cell painting and convolutional neural networks to recognize cell types in dense and mixed cultures with high fidelity. We have benchmarked our approach using pure and mixed cultures of neuroblastoma and astrocytoma cell lines and attained a classification accuracy above 96%. Through iterative data erosion we found that inputs containing the nuclear region of interest and its close environment, allow achieving equally high classification accuracy as inputs containing the whole cell for semi-confluent cultures and preserved prediction accuracy even in very dense cultures. We then applied this regionally restricted cell profiling approach to evaluate the differentiation status of iPSC-derived neural cultures, by determining the ratio of postmitotic neurons and neural progenitors. We found that the cell-based prediction significantly outperformed an approach in which the time in culture was used as classification criterion (96% *vs.* 86%, resp.). In mixed iPSC-derived neuronal cultures, microglia could be unequivocally discriminated from neurons, regardless of their reactivity state. A tiered strategy, allowed for discriminating microglial cell states as well, albeit with lower accuracy. Thus, morphological single cell profiling provides a means to quantify cell composition in complex mixed neural cultures and holds promise for use in quality control of iPSC-derived cell culture models.

## Introduction

Modelling the complexity of the human brain and its (dys)function has proven to be notoriously challenging. This is due to its intricate wiring, the cellular heterogeneity and species-specific differences^1^. To increase the translational value of neuroscientific research, there is a need for physiologically relevant human models. With the advent of human induced pluripotent stem cell (iPSC) technology, it has become possible to generate a wealth of brain-resident cell types including neurons^2^, astrocytes^3^, microglia^4^, oligodendrocytes^5^ and endothelial cells^6^, allowing the study of complex polygenic pathologies that cannot be modelled in animals, opening avenues to precision pharmacology^7,8^. Furthermore, the ability of iPSC to develop into organoids renders them attractive for studying multi-cellular interactions in a 3D context that is closer to the *in vivo* situation^9^. However, genetic drift, clonal and patient heterogeneity cause variability in reprogramming and differentiation efficiency^10,11^. The differentiation outcome is further strongly influenced by variations in protocol^12^. This can significantly impact experimental outcomes, leading to inconsistent and potentially misleading results and consequently, it hinders the use of iPSC-derived cell systems in systematic drug screening or cell therapy pipelines. This is particularly true for iPSC-derived neural cultures, as their composition, purity and maturity directly affect gene expression and functional activity, which is essential for modelling neurological conditions^13,14^. Thus, from a preclinical perspective, there is the need for a fast and cost-effective quality control (QC) approach to increase experimental reproducibility and cell type specificity^15^. From a clinical perspective in turn, robust QC is required for safety and regulatory compliance (e.g., for cell therapy). This need for improved standardization is underscored by large-scale collaborative efforts such as the International Stem Cell Banking Initiative^16^, which focusses on clinical quality attributes and provides recommendations for iPSC validation testing for use as cellular therapeutics, or the CorEuStem network, aiming to harmonize iPSC practices across core facilities in Europe.

Current culture validation methods include (combinations of) sequencing, flow cytometry and immunocytochemistry^15,17^. These methods are often low in throughput, costly and/or destructive. Thence, we set out to develop a method for evaluating the composition of such cultures based on high-content imaging, which is fast, affordable and scalable. The primary goal was to facilitate cell type identification in dense cultures, while ensuring compatibility with subsequent immunocytochemistry or molecular assays for further biological inquiries. To this end, we employed the Cell Painting (CP)^18,19^ method, which is based on labelling cells with simple organic dyes and analysing the resulting phenotypes. CP has proven to be a powerful and generic method for predicting the mode-of-action of pharmacological or genetic perturbations, and this sheerly using a cell morphology readout^20–22^. Thus far, CP has primarily been utilized to predict conditions associated with pharmacological treatments or genetic modifications using images as input. For example, CP has allowed predicting patient diversity in lung adenocarcinoma-associated somatic variants^23^. or genetic variation across donors in iPSC cultures ^24^. Methods such as PhenoRipper^25^ and CP-CHARM^26^ use whole-image features for classification, which circumvent the difficulty of cell segmentation allow classification of images with cells of similar phenotype or class ^27^. However, they do not consider differences in cell density and disregard the inter-cellular heterogeneity within the field of view. Therefore, we explored the amenability of CP to predict individual cell types in dense and mixed cultures. By combining deep learning for cell segmentation and classification, we established an approach that allows recognizing cell types with high accuracy, even in very dense cultures. We employed the approach to evaluate the composition of iPSC-derived neural cultures as they are often quite dense and composed of heterogenous cell types. Varying the density, composition, and cell state allowed us to benchmark the discriminatory potential of cell-based profiling.

## Results

### Neural cell lines have a unique morphotextural fingerprint

Several groups have demonstrated that morphological cell profiling can be used to discriminate pharmacogenomic perturbations based on phenotypic similarity^20–22^. We asked whether a similar approach could be leveraged to unequivocally distinguish individual cell types as well. To this end, we implemented a version of CP based on 4-channel confocal imaging (**Fig. 1A**) and first applied it to monocultures of two neural cell lines from a different lineage, namely astrocyte-derived 1321N1 astrocytoma cells and neural crest-derived SH-SY5Y neuroblastoma cells. First, we explored whether traditional morphotextural feature extraction provided sufficient distinctive power. The features were calculated for each channel in three regions of interest, namely the nucleus, cytoplasm and whole cell. They describe shape, intensity and texture features of each ROI (**Table 5**). Representation of the resulting standardized feature set in UMAP space revealed a clear separation of both cell types along the first UMAP dimension. Clustering of instances was less pronounced after principal component analysis (**Suppl. Fig. 1A**), but UMAP better preserves local and global data structure ^28^. Despite some separation of replicates across the second UMAP dimension, the absence of clear replicate clusters showed that the morphological differences between cell types were consistent across biological replicates (**Fig. 1B**). When projecting individual features onto the UMAP space, we found that both texture (*e.g.,* Nucleus channel 3 energy) and shape (*e.g.,* Cellular Area) metrics contributed to the cell type separation (**Fig. 1C**). The contribution of intensity-related features (*e.g.,* Channel 3 Intensity) to cell type separation was less pronounced as they were more correlated with the biological replicate. Thus, we conclude that cell types can be separated across replicates based on a morphotextural fingerprint.

**Figure 1.**
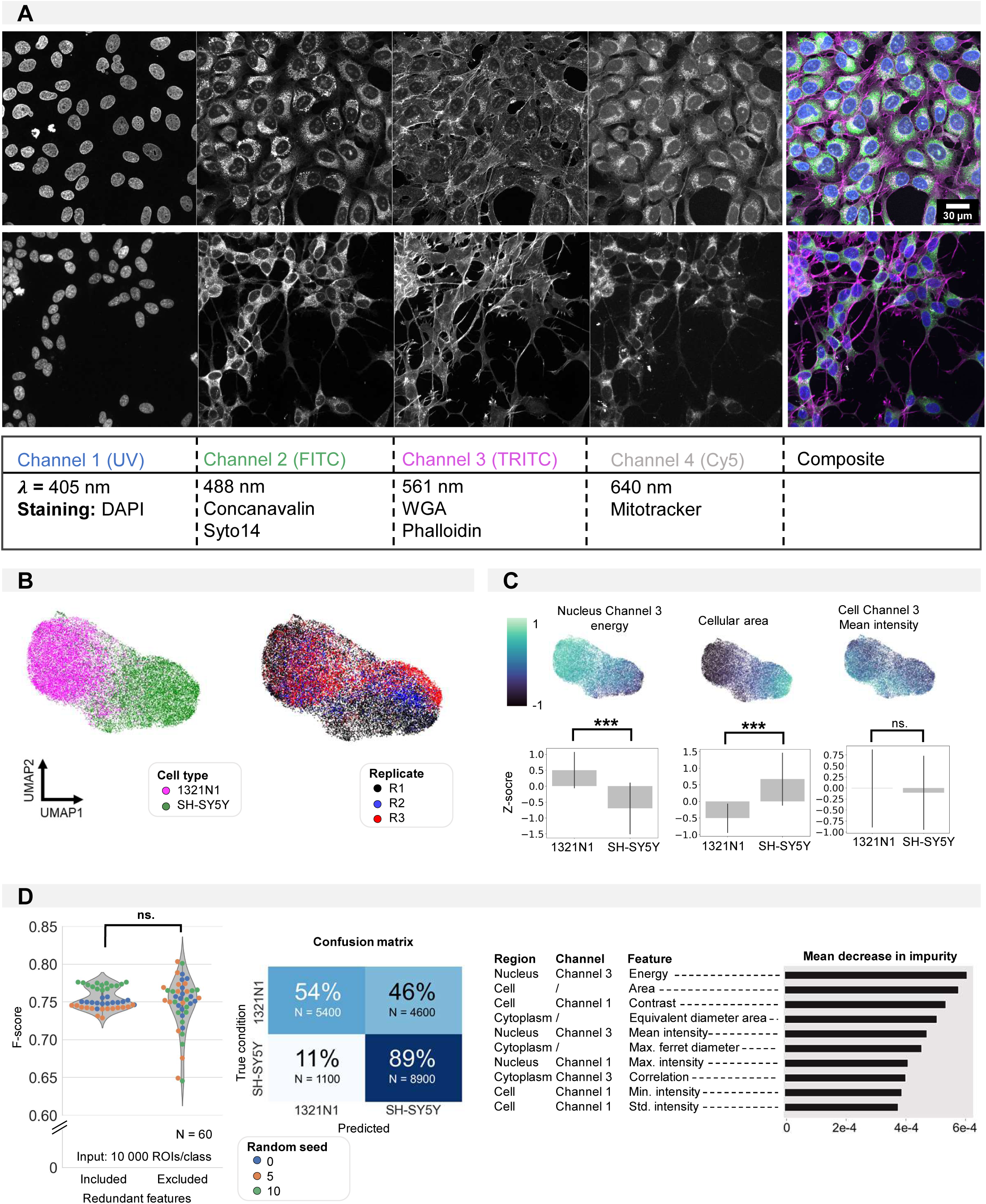
Shallow classification using manually defined features. (A) Image overview of 1321N1 (top) and SH-SY5Y (bottom) cells after CP staining acquired at higher magnification (Plan Apo VC 60xA WI DIC N2, NA = 1,20). Scale bar is 30µm. The channel, wavelength and dye combination used is listed in the table below the figure. This color code and channel numbering is used consistently across all figures. (B) UMAP dimensionality reduction using handcrafted features. Each dot represents a single cell. The color reflects either cell type condition (left) or replicate (right). This shows UMAP clustering is a result of cell type differences and not variability between replicates. (C) Feature importance deducted from the UMAP (feature maps). Each dot represents a single cell. Three exemplary feature maps are highlighted alongside the quantification per cell type. These feature representations help understanding the morphological features that underly the cluster separation in UMAP. (D) Random Forest classification performance on the manually defined feature dataframe with and without exclusion of redundant features. Average confusion matrix (with redundant features) and Mean Decrease in Impurity (reflecting how often this feature is used in decision tree splits across all random forest trees). All features used in the UMAP are used for RF building. Each dot in the violinplot represents the F-score of one classifier (model initialization, N = 30). Classifiers were trained 10x with 3 different random seeds.

### CNN outperforms random forest in cell type classification

To evaluate whether the morphotextural fingerprint could be used to predict cell type, we performed Random Forest (RF) classification using the full feature dataset, using different seeds for splitting up the data into training and validation sets. This resulted in a rather poor accuracy (F-score: 0,75±0,01), mainly caused by the significant (46%) misclassification of 1321N1 cells (**Fig. 1D**). The imbalance in recall and precision was surprising given the clear separation of cell populations in UMAP space using the same feature matrix. When inspecting the main contributions to the RF classifier using the mean decrease in impurity, we found very similar features as highlighted in UMAP space to add to the discrimination (**Fig. 1C**). Where the most important features (*e.g.,* cellular area, channel 1 contrast) showed the expected gradient along the first UMAP direction, lower ranked parameters (e.g., cellular channel 3 mean intensity) had no contribution to UMAP separation (**Fig. 1C**). Reducing noise by removing redundant features (correlation > 0.95) could not ameliorate RF classification performance, which may be due to the documented bias in feature selection for node splitting in high-dimensional data^29^. This result drove us to evaluate a different classification approach based on a ResNet^30^ convolutional neuronal network (CNN). Here, we no longer relied on the extraction of “hand-crafted” features from segmented cell objects for training the shallow RF classifier. Instead, we used isotropic image crops of 60µm centred on individual cell centroids and blanked for their surroundings as input for the CNN (**Fig. 2A**). Using this approach, we found a significantly higher prediction performance with an F-score of 0,96±0,01, with a much more balanced recall and precision (**Fig. 2B**). The UMAP space built from the CNN feature embeddings showed a clear cell type separation (**Suppl. Fig. 1B**). Even with only 100 training instances per class, the CNN outperformed RF, but optimal performance was attained with 5000 training instances (**Fig. 2C**). Both RF and CNN models trained on a combination of three biological replicates (“Mixed reps.”) performed similar as models that were trained on only a single replicate (“Single reps.”). However, the latter does show less variability between different model initializations (**Fig. 2D**). However, a model trained on a dataset containing multiple replicates (“Mixed reps.”) outperformed CNN models that were only trained on a dataset with less variability (“Single rep.”) when predicting instances of an independent unseen replicate (**Fig. 2E**). This emphasizes the need for including sufficient variation in the training set. Although much more performant, CNN classification does not allow direct retrieval of intuitive features, which complicates model interpretation. To gain a visual understanding of image information contributing most to the classification we therefore resorted to Gradient-weighted Class Activation Mapping (Grad-CAM)^31^. This revealed that the attention of the CNN was mainly focused on cell borders (edge information), nuclear and nucleolar signals (**Fig. 2F**). When scrutinizing CNN misclassifications, we found that these are mainly caused by faulty cell detection (*e.g.,* oversegmentation of cell ramifications), unhealthy/dead cells, debris or visibly aberrant cell shapes (**Suppl. Fig. 1C**) - errors not necessarily attributed to the CNN.

**Figure 2.**
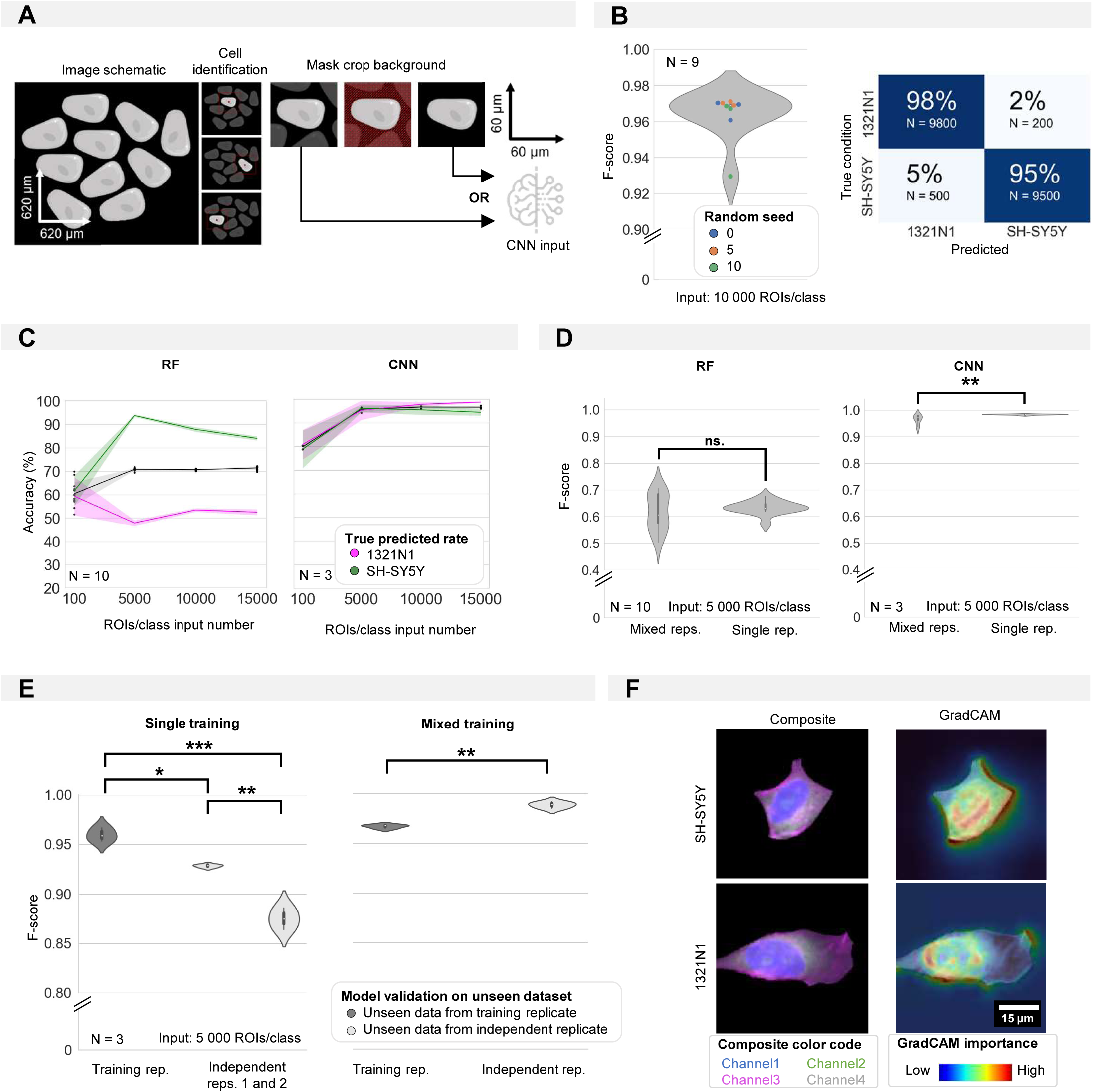
Convolutional neural network classification of monoculture conditions. (A) Schematic of image pre-processing for CNN classification. (B) CNN accuracy and confusion matrix on cell type classification in monocultures. Each dot in the violinplot represents the F-score of one classifier (model initializations, N = 9). (C) Shallow vs. deep learning accuracy for a varying number of input instances (ROIs). Each dot in the violinplot represents the F-score of one classifier (model initialization, N). The ribbon represents the standard deviation on the classifiers. (D) The impact of experimental variability in the training dataset on model performance for shallow vs. deep learning. Classifiers were trained on either 1 (single rep.) or multiple (mixed reps.) replicates (where each replicate consists of N new biological experiment). Mann-Whitney-U-test, p-values resp. 0.7304 and 0.000197. Each violinplot represents the F-score of one classifiers (model initializations). (E) The performance of deep learning classifiers trained in panel D on single replicates (low variability) or mixed replicates (high variability) on unseen images from either the training replicate (cross-validation) or an independent replicate (independent testing) (where each replicate consists of a new biological experiment). Kruskal-Wallis test on single training condition, p-value of 0,026 with post-hoc Tukey test. Mann-Whitney-U-test on mixed training, p-value of 7,47e-6. Each violinplot represents the F-score of one classifier (model initialization, N = 3). (F) Images of example inputs given to the CNN. The composite image contains an overlay of all CP channels (left). The GradCAM image overlays the GradCAM heatmap on top of the composite image, highlighting the most important regions according to the CNN (right). One example is given per cell type.

To shed light on the contribution of individual markers to the classification, we eroded the input to single channel images. For all cases, the prediction performance was below or equal to 85,0% suggesting that no single channel contains all relevant information (**Fig. 3A**). Combinations of two or three channels could not match the prediction accuracy of the full four-channel image either. Thus, we concluded that all channels contribute to the successful classification. As the CNN directly uses image crops as input, the image quality will determine the classification performance. Therefore, we assessed the impact of resolution and signal to noise ratio (SNR). We simulated the effect of decreasing spatial resolution through progressive pixel binning (from 192 pixels (original, pixel size 0.3µm) to 9 pixels (pixel size 6 µm)). For each iteration, 3 CNN models were trained and evaluated. A reduction in pixel size from 0.3µm to 0.6µm did not result in a significantly lower prediction performance, but decreasing the spatial resolution further caused a progressive decrease in F-score (**Fig. 3B**). In a similar manner, we tested the impact of decreasing SNR (by increasing the level of Gaussian noise) on classification accuracy. Starting from an original SNR value of 20,05 dB, we found that the F-score started to decrease when lowering the SNR below 14,27 dB (**Fig. 3C**).

**Figure 3.**
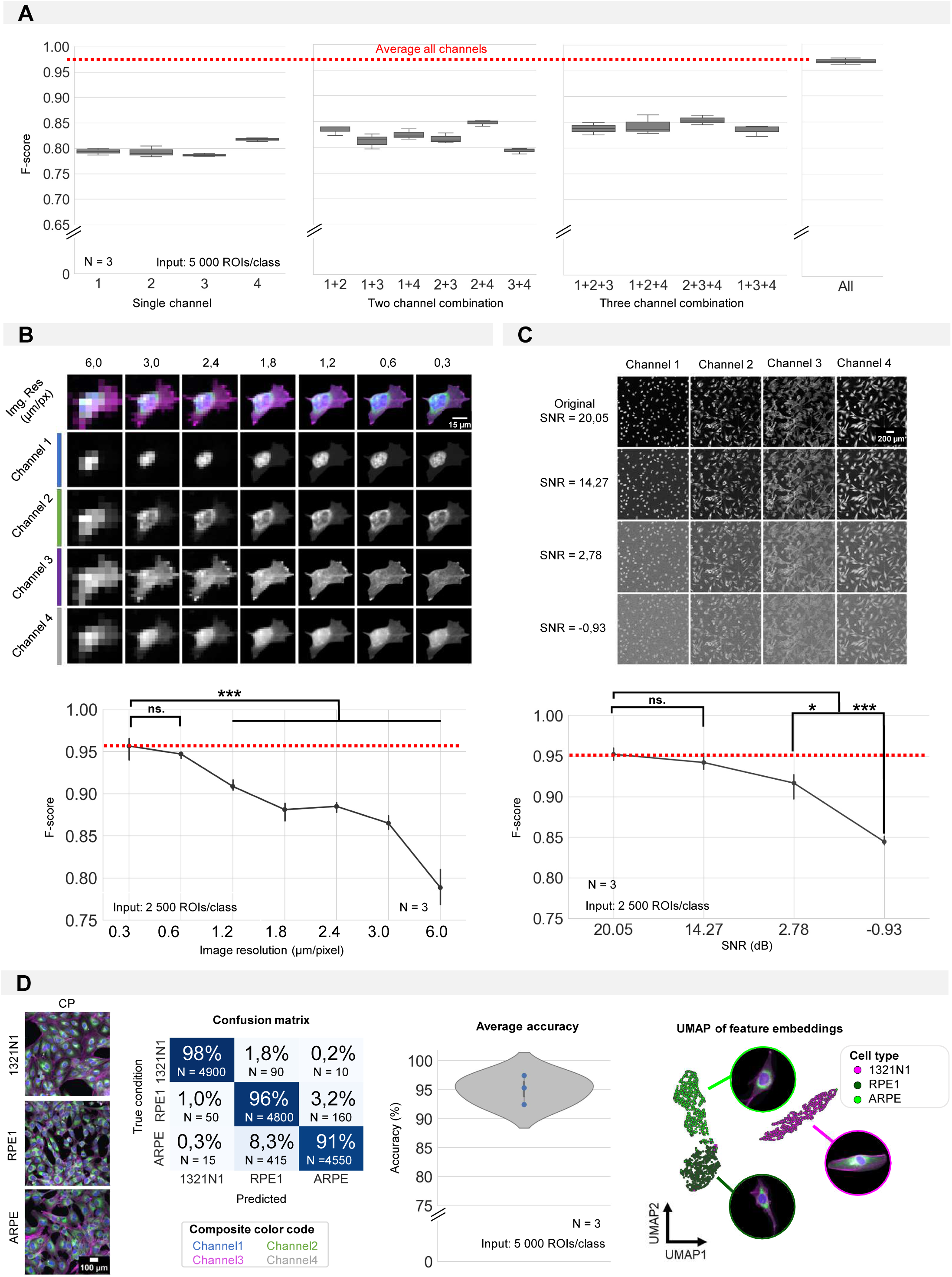
Required input quality for cell type classification. (A) Performance of the CNN when only a selection of image channels is given as training input. Each boxplot represents the F-score of one classifier (model initialization, N = 3). Channel numbering is in accordance with fig. 1A. (1 = DAPI; 2 = FITC; 3 = Cy3; 4 = Cy5) (B) Simulation of the effect of increased pixel size (reduced spatial resolution) on the classification performance. Each point is the average of 3 CNN model initializations (N), with bars indicating the standard deviation between models. Red line indicates the average F-score of the original crops. (C) Simulation of the effect of added gaussian noise (reduced signal-to-noise ratio) on the classification performance. Each point is the average of 3 CNN model initializations, with bars indicating the standard deviation between models. Red line indicates the average F-score of the original images. Statistics were performed using a Kruskal-Wallis test with post-hoc Tukey test. (D) CNN performance on 1321N1, RPE1 and ARPE cells. Dots represent different model initializations (N = 3).

We observed that even with low input image quality, the CNN performance still retained approximately 80% accuracy. We attribute this to the predictive power of the nuclear size, which is a dominant discriminating feature between the cells (**Suppl. Fig. 2**). Given the relatively overt phenotypic difference between 1321N1 and SH-SY5Y cells, we asked whether the CNN could also discriminate more subtle phenotypes. To this end, we used 1321N1 cells and two related retinal pigment epithelial cell lines, hTERT-immortalized RPE1 and the spontaneously immortalized variant ARPE (**Fig. 3D**). The CNN discriminated all three cell lines with high accuracy of 95,06±2,51%. Unsupervised UMAP dimensionality reduction using the CNN feature embeddings revealed three clusters of ROIs, with the two RPE lines located closer together (but well separated) in comparison to the morphologically more distinct 1321N1 cell type (**Fig. 3D**). Together this work illustrates that a CNN approach can be used to distinguish diverse cell types within a given image quality window.

### Nucleocentric predictions remain accurate regardless of culture density

Morphological profiling relies on accurate cell detection. This may become difficult in dense cultures such as iPSC-cultures, clustered cells tissues and tissue-mimics. Having established a method to distinguish cell types with high accuracy, we next asked how robust the classification was to increasing cell density. To this end, we acquired images of 1321N1 and SH-SY5Y monocultures, grown at densities ranging from 0 to 100% confluency (**Fig. 4A**). Based on the nuclear count, we binned individual fields into 6 density classes (0-20%, 20-40%, 40-60%, 60- 80%, 80-95%, 95-100%) and trained a CNN with equal sampling of cell numbers per density class to avoid bias. No decrease in accuracy was observed until the culture density reached 80% confluency. Only for very high densities (95-100%), we found a significant decrease in the prediction accuracy (F-score: 0,92±0,05) (**Fig. 4B**). We reasoned that under these conditions cell shape would be predominantly determined by neighbouring cells and cell segmentation performance would decrease (**Suppl. Fig. 1D**). Nuclei are less malleable and even in dense cultures remain largely separated, allowing their robust segmentation and avoiding CNN misclassifications resulting from segmentation errors. Hence, we asked whether using the nuclear ROI as input would improve classification performance at high densities. However, despite the relatively high average F-score of 0,91±0,05 (**Fig. 4C**), the performance was consistently lower than whole cell ROI across the density range and the performance still decreased with full confluency. To understand these results, we inspected the GradCAM output for these predictions, and found that an important part of the attention of the CNN is diverted to the background (**Fig. 4D, Suppl. Fig. 3**). We interpret this result as the CNN using the background as a proxy for nuclear size. To rule out bias by setting the background to zero in nuclear crops, the CNN was also trained on the same crops with randomly speckled background, but despite a shift in attention to the nucleus, similar prediction performance was attained (**Suppl. Fig. 2**). The classification performance was not biased by segmentation errors (**Suppl. Fig. 1E**) but may be influenced by the fact that nuclear area is affected by culture density (**Suppl. Fig. 1F**). In highly dense cultures, the dynamic range of the nuclear size decreases as all nuclei become more compact and the nuclear size range decreases (**Suppl. Fig. 1F**). Thence, we tested an intermediate condition which exploits the more robust nuclear segmentation but also includes part of the (sub-)cellular local surrounding information as input (**Suppl. Fig. 1G**). To identify the optimal patch size, we varied the size of a square box centred around the nuclear centroid from 0.6 to 150 µm (**Fig. 4E**). Within a range of 12-18µm, we found a maximal F-score of 0,96±0,02. While the F-score remained relatively high with increasing patch sizes up to 60µm, increasing the patch size above 42µm decreased the confidence of the prediction as evidenced by a lower precision and increased variability on the prediction results (**Fig. 4E**). Hence, for further experiments, a nucleocentric patch size of 18µm was used. GradCAM images revealed that this latter approach led the CNN to focus on perinuclear structures (**Fig. 4D and Suppl. Fig. 3)**. Interestingly, when using this nucleocentric approach, the prediction performance was maintained at almost confluent cell densities in contrast with whole cell approaches (**Fig. 4B**). Thus, we conclude that using a nucleocentric region as input for the CNN could be a valuable strategy for accurate cell type identification in dense cultures and that this method is robust to a wide range of patch sizes.

**Figure 4.**
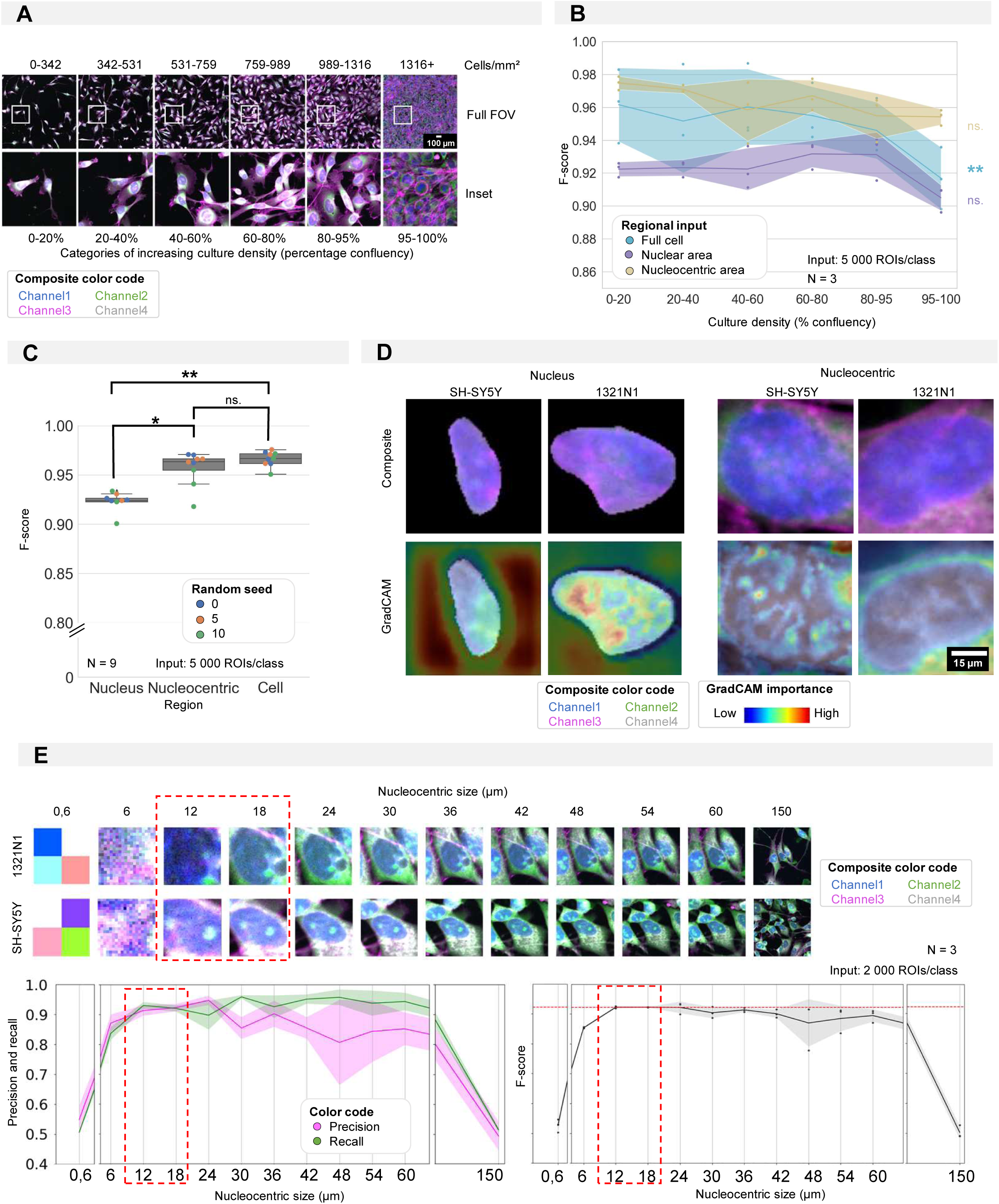
Required regional input for accurate CNN training in cultures of high density. (A) Selected images and insets for increasing culture density with the density categories used for CNN training. (B) Results of 3 CNN models trained using different regional input (the full cell, only nucleus or nucleocentric area) and evaluated on data subsets with increasing density. Each dot represents the F-score of one classifier (model initialization, N = 3) tested on a density subset. Classifiers were trained with the same random seed. Ribbon represents the standard deviation. (C) CNN performance (F-score) of CNN models using different regional inputs (full cell, only nucleus or nucleocentric area). Each boxplot represents 3 model initializations for 3 different random seeds (N = 9). (D) Images of example inputs for both the nuclear and nucleocentric region. The composite image contains an overlay of all CP channels (top). The GradCAM image overlays the GradCAM heatmap on top of the composite image, highlighting the most important regions according to the CNN (bottom). One example is given per cell type. (E) Systematic in- and decrease (default of 18µm used in previous panels) of the patch size surrounding the nuclear centroid used to determine the nucleocentric area. Each dot represents the results of one classifier (model initialization, N = 3). Ribbon represents the standard deviation. The analysis was performed using a mixed culture dataset of 1321N1 and SH-SY5Y cells (Fig. 5).

### Cell prediction remains accurate in mixed cell culture conditions

Although a very high classification accuracy was obtained with nucleocentric CNN predictions, both training and testing were performed with input images drawn from monocultures. As our goal was to allow cell type prediction in complex, heterogeneous cultures, we next switched to a more faithful situation in which we co-cultured both cell types. Ground-truth for these predictions was generated by either performing post-hoc immunofluorescence (IF) with cell-type specific antibodies (CD44 for 1321N1 and TUBB3 for SH-SY5Y cells), or by differential replication labelling (*i.e.,* by incubating the two cell types with EdU and BrdU respectively, prior to mixing them) after dye quenching^32^ (**Fig. 5A**). Replication labelling proved significantly more successful for binary ground truth classification (through intensity thresholding) than IF labelling for the cell lines we used (**Fig. 4B**). When training a CNN to recognize the cell types using these markers as ground truth, we found that the prediction accuracy on the left-out dataset of the co-culture (Co2Co) was almost as high as when a model was trained and tested on monocultures (Mono2Mono) (**Fig. 5C**).

**Figure 5.**
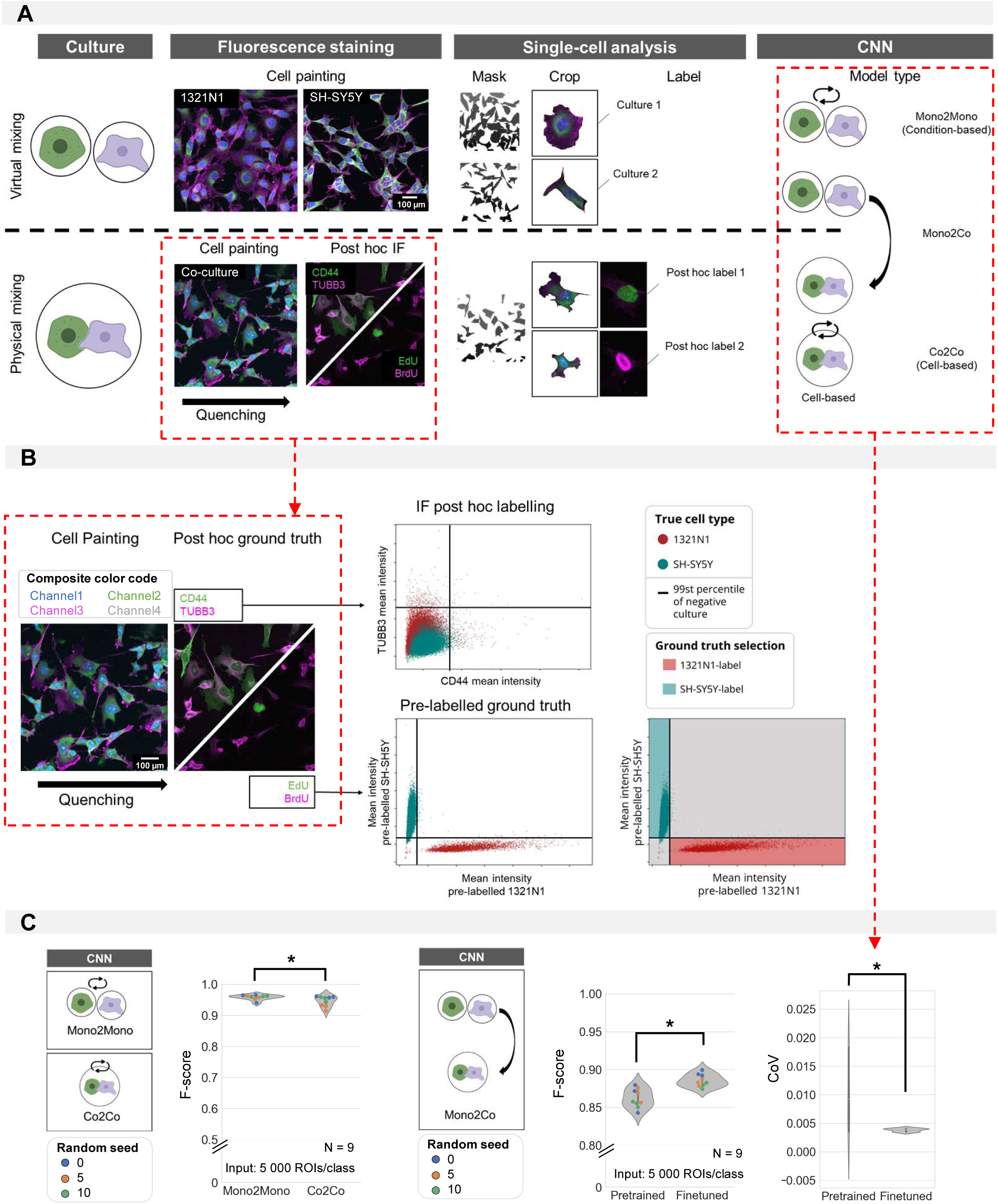
Cell type prediction in mixed cultures by post-hoc ground truth identification. (A) Schematic overview of virtual vs. physically mixed cultures and the subsequent CNN models. For virtual mixing, individual cell types arise from distinct cultures, the cell type label was assigned based on the culture condition. For physical mixing, the true phenotype of each crop was determined by intensity thresholding after post-hoc staining. Three model types are defined: Mono2Mono (culture-based label), Co2Co (cell-based label) and Mono2Co (trained with culture-based label and tested on cell-based label). (B) Determining ground-truth phenotype by intensity thresholding on IF (top) or pre-label (bottom). The threshold was determined on the intensity of monocultures. Only the pre-labelled ground-truth resulted in good cell type separation by thresholding for 1321N1 vs. SH-SY5Y cells. (C) Mono2Mono (culture-based ground truth) vs. Co2Co (cell-based ground truth) models for cell type classification. Analysis performed with full cell segmentation. Mann-Whitney-U-test p-value 0,0027. Monoculture-trained models were tested on mixed cultures. Pretrained models were trained on independent biological replicates. These could be finetuned by additional training on monoculture images from the same replicate as the coculture. This was shown to reduce the variation between model iterations (Median performance: Mann-Whitney-U-test, p-value 0,0015; Coefficient of variation: Mann-Whitney-U-test, p-value 3,48e-4). Each dot in the violinplots represents the F-score of one classifier (model initialization, N = 9). Classifiers were trained 3x with 3 different random seeds.

We then tested whether it was possible to train a classifier based on monocultures only, for cell type prediction of cells in co-culture (Mono2Co). This resulted in an F-score of 0,86±0,01%. This drop in performance may be caused by the effect on cell phenotype exerted by the presence of other cell types in mixed cultures which is not captured in monocultures. As the images from monocultures and co-cultures were obtained from different plates, we suspected inter-replicate variability in culture and staining procedures to contribute in part to the lesser performance. Therefore, we tested whether we could improve the performance of the CNN by including monocultures from the same plate as the co-cultures. This finetuning indeed improved the average performance to 0,88±0,01%, and more importantly, it significantly reduced the variability (coefficient of variation) of the predictions making it more reproducible (**Fig. 5D**). Thus, while not yet reaching the same accuracy, it proves that it is possible to establish a model that recognizes cell types in co-cultures that is solely trained on monocultures.

### Cell type profiling can be applied to stage iPSC-derived neuronal cultures

iPSC-derived neuronal cell cultures suffer from significant inter- and intra-batch variability and could benefit from an efficient quality control^14,33^. Thence, we applied our nucleocentric phenotyping pipeline to stage the maturity of a neuronal cell culture based on its cell type composition. Using a guided protocol^2^, two differentiation stages were simulated: a primed stage, where most cells are assumed to be cycling neural progenitors (NPCs), and a differentiated stage where most cells are post-mitotic neurons (**Fig. 6A**). The two cell types were discriminated by post-hoc IF labelling for the cell cycle marker Ki67 (NPC) and the microtubule marker ß-III-tubulin (TUBB3, neurons) (**Fig. 6B**). Not all cells in the CP image could be assigned with a ground truth label due to cell detachment upon re-staining or sheer absence of the tested marker. Since no single monoculture consists of 100% of either cell type, we applied gates to retain only those cells that show unequivocal staining for either one of both markers. Based on these gates, ROIs were either classified as neuron (Ki67-/TUBB3+), NPC (Ki67+/TUBB3-) or undefined (outside of gating boundaries). We assume the latter category to represent transitioning cells in intermediate stages of neural development, un- or dedifferentiated iPSCs. This gating strategy resulted in a fractional abundance of neurons vs. total (neurons + NPC) of 36,4 % in the primed condition and 80,0% in the differentiated condition (**Fig. 6C**). In a first attempt to classify cells within the guided culture, we trained a CNN on individual cell inputs using the culture condition (primed or differentiated) as ground truth, which given the heterogeneity, can be considered as a weak label. This resulted in a low F-score of 0,86±0,01%. When we used the cell-level IF ground truth labels instead, we obtained a classification performance of 0,96±0,00 %. For comparison, a shallow learner (RF), showed a significantly lower F-score of 0,87±0,02 (**Fig. 6D**). Applying this cell-based (as opposed to condition-based) CNN to the two culture conditions resulted in predicted fractional abundance of neurons to NPC of 40,5% in images of the primed condition and 74,2% in images of the differentiated condition – aligning well with the manually defined ratios (**Fig. 6E**). Both supervised and unsupervised UMAP dimensionality reduction on the feature embeddings of the cell-based classifier revealed a clear clustering of both phenotypes suggesting that the CNN captures the differences in morphotextural fingerprint between neurons and NPCs well (**Fig. 6F**). We then went on to evaluate the established cell-based classifier to a primed neuronal cell culture undergoing gradual spontaneous differentiation after dual SMAD inhibition^34^. We examined cell cultures at 13, 30, 60 and 90 days *in vitro* (DIV) after the start of the differentiation process and visually confirmed a gradual change in neural maturity. This is evidenced by a gradually smaller cell body and long, thin ramifications (**Fig. 6G**). The cell-based CNN model trained on the guided differentiation dataset tested on the spontaneous cultures, confirmed a shift in the (slower) neuron-to-NPC fractional abundance with increasing time in culture (**Fig. 6H**).

**Figure 6.**
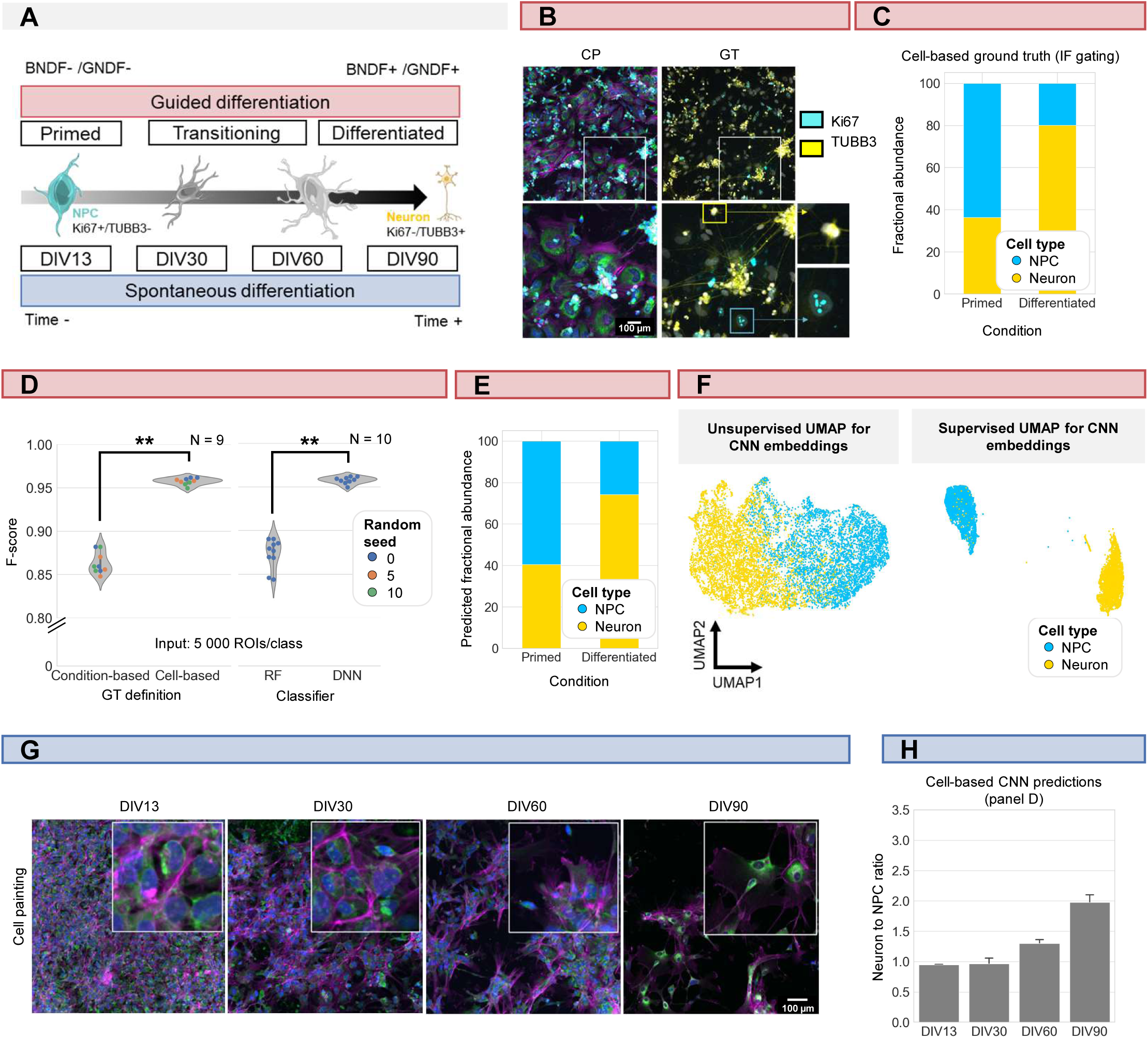
iPSC-derived differentiation staging using morphology profiling. (A) Schematic overview of guided vs. spontaneous neural differentiation. DIV = days in vitro. Selected timepoints for analysis of the spontaneously differentiation culture were 13, 30, 60 and 90 days from the start of differentiation of iPSCs. (B) Representative images of morphological staining (color code as defined in fig. 1A) and post-hoc IF of primed and differentiated iPSC-derived cultures (guided differentiation). Ground-truth determination is performed using TUBB3 (for mature neurons) and Ki67 (mitotic progenitors). (C) Fraction of neurons vs. NPC cells in the primed vs. differentiated condition as determined by IF staining. Upon guided differentiation, the fraction of neurons increased. (D) left: CNN performance when classifying neurons (Ki67-/TUBB3+) vs. NPC (Ki67+/TUBB3-) cells using either a condition-based or cell-type based ground truth. Each dot in the violinplots represents the F-score of one classifier (model initialization). Classifiers were trained with different random seeds. Mann-Whitney-U-test, p-value 4,04e-4. Right: comparison of CNN vs. RF performance. Mann-Whitney-U-test, p-value 2,78e-4. (E) Fractional abundance of predicted cell phenotypes (NPC vs. neurons) in primed vs. differentiated culture conditions using the cell-based CNN. (F) Unsupervised and supervised UMAP of the cell-based CNN feature embeddings. Plot color coded by cell type. Points represent individual cells. (G) Representative images of spontaneously differentiating neural cultures. Color code as defined in fig. 1A. (H) Prediction of differentiation status using the cell-based CNN model trained on guided differentiated culture.

### Cell type identification can be applied to mixed iPSC-derived neuronal cultures regardless of the activation state

Next to NPCs and neurons, iPSC-derived neuronal cultures are often studied in conjunction with other relevant cell types that influence neuronal connectivity and homeostasis such as astrocytes and microglia^13^. Therefore, we tested whether the cell-based approach could be extended to these cell types as well. We generated monocultures of iPSC-derived astrocytes, neurons, and microglia from the same parental iPSC line (**Fig. 7A**) and trained a nucleocentric CNN to identify each cell type. This led to a prediction accuracy of 96,81±0,95% (**Fig. 7A**). Recognizing that these cell types are visually morphologically distinct, we tested the robustness of the model to experimental perturbations in cell state (**Fig. 7B**). To this end, we induced reactivity in the iPSC-derived microglial culture by LPS treatment. A CNN trained on monocultures of neurons, unchallenged or LPS-treated microglia showed high accuracy in differentiating neurons from microglia (98% accurate), although the difference between reactive and resting-state microglia in this tripartite model proved more challenging (71% and 90% accuracy, resp.). However, when implementing a tiered approach, in which after neuron-microglia recognition, a second model was tasked with the classification of reactive vs. resting state microglia without the presence of neurons, an F-score of 0,92±0,00 was obtained. Repeating those experiments in mixed cultures of neurons and microglia (using Tubb3, resp. Iba1 as ground-truth IF markers), yielded an F-score of 0,98±0,01 for the classification of neurons vs. microglia. Again for comparison, CNN outperformed the classical RF approach (0,86±0,03) (**Fig. 7C**). In the presence of LPS, a CNN model yielded a high F-score of 0,97±0,01 (**Fig. 7C**). This implies that the shift in cell state does not significantly affect the CNN’s ability to distinguish neurons from microglia. As above, a separate CNN was trained to classify within the microglial subpopulation the LPS-treated (reactive) from the unchallenged (resting-state) microglia. Yet, this phenotype proved more challenging for the CNN to predict, as evidenced by an F-score of 0,80±0,01 (**Fig. 7C**). Based on these results, we conclude that nucleocentric phenotyping can be used to gauge the cell type composition of iPSC-derived cultures. Distinct cell types show sufficient morphological differences to allow highly accurate classification. Different cell states of individual cell types can still be recognized albeit with lower performance.

**Figure 7.**
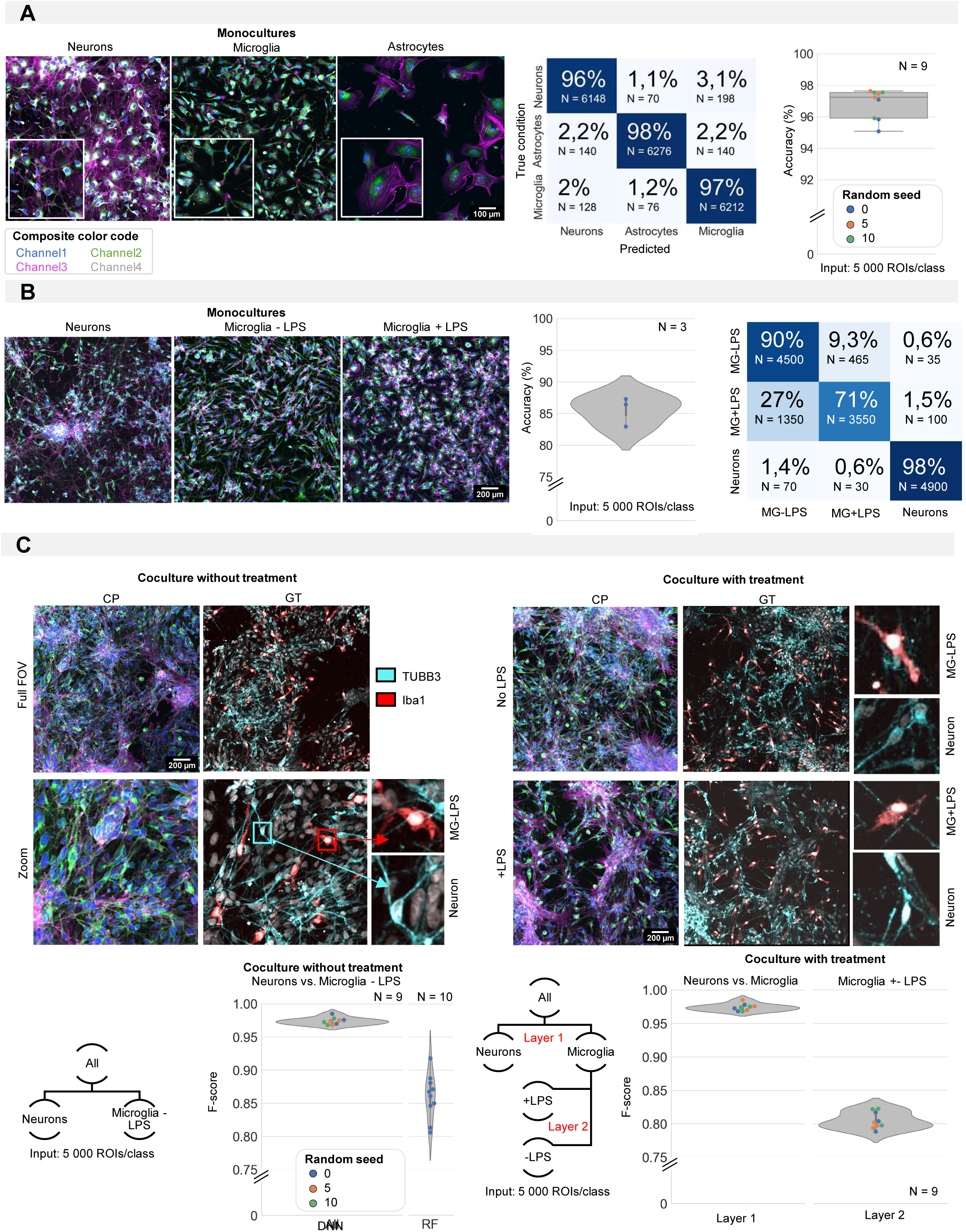
iPSC cell type identification using morphology profiling. (A) Representative images of iPSC-derived neurons, astrocytes and microglia in monoculture with morphological staining. Color code as defined in fig. 1A. Prediction accuracy of a CNN trained to classify monocultures of iPSC-derived astrocytes, microglia and neurons with confusion matrix (average of all models). Each dot in the boxplot represents the F-score of one classifier (model initialization, N = 9). Classifiers were trained 3x with 3 different random seeds. (B) Representative images of monocultures of iPSC-derived neurons and microglia treated with LPS or control. Color code as defined in fig. 1A. Prediction accuracy and confusion matrix (average of all models) are given. Each dot in the violinplot represents the F-score of one classifier (model initialization, N = 3). (C) Representative images of a mixed culture of iPSC-derived microglia and neurons. Ground-truth identification was performed using IF. Each dot in the violinplot represents the F- score of one classifier (model initialization). Classifiers were trained 3 different random seeds. Results of the CNN are compared to shallow learning (RF). The same analysis was performed for mixed cultures of neurons and microglia with LPS treatment or control. A layered approach was used where first the neurons were separated from the microglia before classifying treated vs. non-treated microglia. Each dot in the violinplot represents the F-score of one classifier (model initialization, N = 9). Classifiers were trained 3x with 3 different random seeds.

## Discussion

iPSC-derived cell cultures have the potential to improve translatability of biomedical research but represent a challenge due to their variability and multicellular composition. With our work, we have developed a method to identify neural cell types in mixed cultures to aid in their validation and application in routine screening settings. We first benchmarked our approach using neural cell lines and found that a CNN-based approach outperforms shallow learning (RF) based on handcrafted feature extraction, even with a limited number of input images. This aligns well with recent data showing CNN-based feature representations outperform shallow techniques for condition-based drug response prediction^35^. To assess the robustness and sensitivity of deep learning for cell type prediction, we evaluated the CNN performance as a function of input data size and quality. Next, we tested the performance of predictions for dense single and multi-cellular cultures. Finally, we assessed its performance in the presence of different cells states.

Although our benchmarking revealed a minimal image resolution of 0,6 µm/pixel and SNR of approximately 10 dB for optimal CNN performance, lower quality images may still yield acceptable performance, especially with the use of advanced image restoration algorithms^36^. We found that all channels contribute to the overall prediction accuracy but that the whole cell is not required to obtain the highest prediction accuracy. In line with earlier studies that highlight the biomarker potential of the nucleus for patient stratification^37^, cell state identification^38^ or drug response^20^, we found that the nuclear ROI as such already carries highly distinctive information for cell type prediction. However, extension to its direct surroundings proved to be the best and most robust input across a range of cell densities. The nucleocentric approach, which is based on more robust nuclear segmentation, reduces segmentation errors whilst still retaining input information from the structures directly surrounding the nucleus. At higher cell density, the whole-cell segmentation becomes more error-prone, while also losing morphological information (**Suppl. Fig. 1D**). When using a patch capturing the nucleus and its close environment (within a relatively large size range) predictions were much more consistent across the cell density range. We assume this is due to the more robust nuclear (vs. cellular) segmentation and the fact that it does not blank the surrounding regions. The latter also buffers for occasional nuclear segmentation errors (*e.g.,* where blebs or parts of the nucleus are left undetected). GradCAM maps on nucleocentric crops highlight specifically those structures surrounding the nucleus (reflecting ER, mitochondria, Golgi) indicating their importance in correct cell classification. It opens possibilities for applying cell profiling in the future to even more complex 3D cell systems such as tissue or organoid slices, where accurate cell segmentation becomes extremely challenging. Indeed, our results indicate that it might be possible to discriminate individual cells in extremely crowded environments. For specific applications such as classification of healthy vs. tumour tissue, binary classification models can prove useful. It has been shown that a dual-task graph neural network can classify epithelioid and sacromatoid tumor cells with cellular resolution for accurate mesothelioma subtype determination^39^. However, the rich diversity of cell types and intermediates present in tissue may complicate classification and demand a more extensive ground truth based on cyclic immunostaining to further refine the predictions. In the further future, it may even be possible to include volumetric information, but this will require optimization of the sample preparation procedures as well.

One crucial realisation of this work is that cell types can be identified in mixed cultures solely using the input from monocultures. This implies that cells retain their salient characteristics even in more complex heterogeneous environments. While for now, predictions are still superior when based on training directly on mixed cultures, we found that the prediction accuracy of monoculture trained models can be increased by employing replicate controls. This suggests that it may become possible to apply or adapt existing models without the need for a cell-level ground truth as provided by post-hoc labelling. This technique could potentially be of use for cultures or tissues where no antibody- or pre-labelling is possible (*e.g.,* no unique IF marker is available, non-replicating cells). To increase the accuracy one could resort to intelligent data augmentation^40^ or transfer learning^41^ strategies. While a ground-truth independent method holds promise for bulk predictions (*e.g.,* for quality control purposes), the use of post-hoc labelling allows building more refined models that can distinguish multiple cell types and/or cell states at once. Especially using cyclic staining or spatial transcriptomics, much richer information content can be gained. With the method presented here, we combine the rich multidimensional information from cyclic IF with morphological information and show that it is possible to predict IF signatures. Multiplexed imaging has previously shown its utility in gaining in-depth information on culture composition and differentiation status in iPSC-derived neural cultures (progenitor, radial glia, astrocytes (immature and mature) and neurons (inhibitory and excitatory)^42^. Similarly, this could be expanded to cell states (apoptosis, DNA damage), albeit with a lesser performance depending on the phenotype^21^. Initial model training requires both CP and multiplexed IF images. After the initial training phase, the IF step can be omitted, and the CNN will predict the phenotype outcomes of the IF with high accuracy on the CP images alone. This significantly reduces experimental costs and time requirements.

Applying the method to iPSC-neuronal cultures revealed its potential to score their differentiation state. Although guided protocols manage to speed up the differentiation process and lead to enhanced culture purity, the neural differentiation process proves to be less than binary, as evidenced by the mixed presence of Ki67+/TUBB3- and Ki67-/TUBB3+ cells. The spontaneous differentiation protocol illustrated the unpredictable nature and late stage of the differentiation process. Many groups highlight the difficulty of reproducible neural differentiation and attribute this to culture conditions, cultivation time and variation in developmental signalling pathways in the source iPSC material^43,44^. Spontaneous neural differentiation has previously been shown to require approximately 80 days before mature neurons arise that can fire action potentials and show neural circuit formation. Although these differentiation processes display a stereotypical temporal sequence^34^, the exact timing and duration vary. This variation negatively affects the statistical power when testing drug interventions and thus prohibits the application of iPSC-culture derivatives in routine drug screening. Current solutions (e.g., immunocytochemistry, flow cytometry, …) are often cost-ineffective, tedious, and incompatible with longitudinal/multimodal interrogation. CP is a much more cost-effective solution and ideally suited for this purpose. Routine CP-based could add confidence to and save costs for the drug discovery pipeline. We have shown that CP can be leveraged to capture the morphological changes associated with neural differentiation. We could reliably predict the gradual transition from NPC to a more differentiated state. We realize that differentiating iPSC cultures are highly heterogenous and are composed of a landscape of transitioning progenitors in various stages of maturity that our binary models currently fail to capture. As a result, we filtered out cells with a ‘dubious’ IF profile (e.g., cells that might be transitioning or are of a different type) as they would negatively affect the model by introducing noise. In future iterations, one could envision defining more refined cell (sub-)types in a population based on richer post-hoc information (e.g., through cyclic immunofluorescence or spatial single cell transcriptomics). While we emphasize the value of identifying fixed states as a fast-track to gather the composition of an iPSC-derived culture, it is equally interesting to focus on the continuum that such cultures represent. To predict the maturation of iPSC-derived neural progenitors to differentiated neurons in a more continuous manner, would demand regression-based approaches ^45^. Pioneering efforts using live-cell imaging and machine learning have allowed predicting gradual cell state transitions, for example in the context of myoblast, adipocyte or osteoblast differentiation ^46,47^. Label-free timelapse imaging has also been shown to aid in the assessment of differentiation without disruption of the sample^46^. Given that many of the cell painting dyes are compatible with live cells, it is conceivable that our approach is amenable to further refining the assessment of iPSC differentiation status in real time.

After testing the CNN performance on heterogenous cultures, we have added an additional layer of complexity by inducing microglial reactivity in a coculture of neurons and microglia. We found that we could still predict the cell type regardless of the treatment. The increased variability within the microglial subpopulation did not impact the CNNs’ ability to discriminate cell types in mixed culture. Furthermore, using the layered approach, the resting-state and reactive subgroups within the microglial population could also be classified, albeit with a lesser prediction accuracy. This could be explained by the fact that microglia grown *in vitro* are not completely homeostatic, even in absence of LPS ^48^, and may require different cell types to adopt a more natural state. Having more faithful mixed cultures with cells in their endogenous states, and an apt tool to recognize both cell type and state, holds promise to enhance the relevance of preclinical screens, increase the accuracy of drug targets and ultimately lead to more precise therapeutic strategies^49^. For now, we have tailored models to the individual datasets, but it is conceivable that a more generalized CNN could be established for multiple culture types. This would obviously demand a much larger dataset to encompass the variability encountered in such models (*e.g.,* various starting iPSC lines, various differentiation protocols). Publicly available datasets (*e.g.,* JUMP-Cell Painting Consortium) can aid in creating an input database containing a large variability (different iPSC lines, different neural differentiation protocols, …), which would ultimately lead to a more robust predictor. Our results showing the prediction accuracy of a guided differentiation model on spontaneously differentiating cultures indicate that the approach can be transferred to other differentiation protocols as well. Inclusion of more input images and variability should thus enable developing a generalist model for other differentiation protocols without the need for ground truth validation and further CNN training.

In conclusion, we have developed a novel application for unbiased morphological profiling by extending its use to complex mixed neural cultures using sequential multispectral imaging and convolutional network-informed cell phenotype classification. We show that the resulting predictors are robust with respect to biological variation and cell culture density. This method holds promise for use in quality control of iPSC cultures to allow their routine use in high-throughput and high-content screening applications.

## Methods

### Cell culture

Cells were cultured at 37 °C and 5% CO2. 1321N1 and SH-SY5Y cell lines were maintained in DMEM-F12 + Glutamax (Gibco, 10565018) supplemented with 10% Fetal Bovine Serum (Gibco, 10500064). Cell seeding prior to imaging was done in 96-well black multiwell plates with #1.5 glass-like polymer coverslip bottom (Cellvis, P96-1.5P). Only the inner 60 wells were used, while the outer wells were filled with PBS-/- to avoid plate effects. Plates were coated with Matrigel (Corning, 734-1440) After seeding, the imaging plate was centrifuged at 100g for 3min.

iPSCs (Sigma Aldrich, iPSC0028 Epithelial-1) were cultured on Matrigel (Corning, 734-1440) in Essential 8 medium (Gibco, A1517001). Upon cell thawing, Rock inhibitor (Y-27632 dichloride, MedChem, HY-10583) was added in 10µM concentration. Subculturing of iPSCs was performed with ReLeSR (Stemcell Technologies, 05872).

Unguided differentiation^34^ of iPS cells to neural progenitor cells (NPCs) was started by subculturing the iPSCs single cell using Tryple Express Enzyme (Life technologies, 12605010) at a density of 10e4 cells/cm² in mTesR1 medium (Stemcell Technologies, 85850) and Rock inhibitor. The following day, differentiation to NPCs was started by dual-SMAD inhibition in neural maintenance medium (1:1 Neurobasal (Life technologies, 21103049):DMEM- F12+Glutamax (Gibco, 10565018), 0.5x Glutamax (Gibco, 35-050-061), 0.5% Mem Non Essential Amino Acids Solution (Gibco, 11140050), 0.5% Sodium Pyruvate (Gibco, 11360070), 50 μM 2-Mercaptoethanol (Gibco, 31350010), 0.025% Human Insulin Solution (Sigma Aldrich, I9278), 0.5X N2 (Gibco, 17502048), B27 (Gibco, 17504044), 50 U/ml Penicillin-Streptomycin (Gibco, 15140122)) supplemented with 1 μM LDN-193189 (Miltenyi, 130-106-540), SB431542 (Tocris, 1614). This dual blockade of SMAD signalling in iPSCs is induces neural differentiation by synergistically causing the loss of pluripotency and push towards neuroectodermal lineage^50^. Daily medium changes were performed for 11 consecutive days. Following neural induction, neural rosettes were selected by STEMdiff Neural Rosette Selection Reagent (Stemcell technologies, 05832). Maintenance of neural progenitor cells was performed in neural maintenance medium with 10 µM bFGF (Peprotech, 100-18C) added after subculturing. Single cell detachment of NPCs during maintenance was performed using Tryple Express Enzyme. Cell seeding prior to imaging is done in 96-well black multiwell plates µCLEAR (Greiner, 655090) coated with Poly-L-ornithine (Sigma-Aldrich, P4957) and laminin (Sigma-Aldrich, L2020). Only the inner 60 wells were used, while the outer wells were filled with PBS-/- to avoid plate effects. After seeding, the imaging plate was centrifuged at 100g for 3min.

Guided iPSC differentiation to neurons was performed according to Bell et al. (2019)^2^. The initial phase of neural induction consisted of 12 days neural induction medium (DMEM- F12+Glutamax (Gibco, 10565018), 1x N2 (Gibco, 17502048), 1x B27 (Gibco, 17504044),1mg/ml BSA (Sigma-Aldrich, A7979), 0.5% Mem Non-Essential Amino Acids Solution (Gibco, 11140050)). Of these 12 days, the first 7 were supplemented with cytokines for dual-SMAD inhibition (1 μM LDN-193189 (Miltenyi, 130-106-540), SB431542 (Tocris, 1614)). Following neural induction, the NPCs were floated in uncoated MW6 culture plates in NPC medium (DMEM-F12+Glutamax (Gibco, 10565018), 1x N2 (Gibco, 17502048), 1x B27 (Gibco, 17504044), 10 µM bFGF (Peprotech, 100-18C), 10 µM EGF (Peprotech, 100-47)). After 4 days, NPC clusters of appropriate size were filtered using a cell strainer ((37 µm cell strainer, Stemcell technologies, 27250) and plated on Matrigel-coated (Corning, 734-1440) MW6 culture plates. NPCs can now be expanded in NPC medium. Guided differentiation into forebrain neurons can be induced by switching to neuronal medium (BrainPhys (STEMCELL Technologies, 05792), 1x N2 (Gibco, 17502048), 1x B27 (Gibco, 17504044), 10 µM BDNF (PeproTech, AF-450-02), 10 µM GDNF (PeproTech, AF-450-02) for 15 days before fixation.

Differentiation of iPSC to microglia^4^ was performed by the formation of embroid bodies (EBs) with 10e3 iPSCs/well in a 96-well U-bottom plate (Corning, 351177) coated with Anti-Adherence Rinsing Solution (Stemcell technologies, 07010) in mTeSR medium supplemented with Rock inhibitor, 50 ng/mL BMP4 (Peprotech, 120-05), 50 ng/mL VEGF (Peprotech, 100-20), 20 ng/mL SCF (Peprotech, 250-03). 75% medium is changed for 4 consecutive days. After mesoderm induction, EBs are transferred to a 6-well plate with 20 EBs/well and placed in macrophage precursor medium (X-vivo15 (Lonza, BE02-060Q), 100 ng/mL M-CSF (Peprotech, 300-25), 25 ng/mL IL-3 (Peprotech, 213-13), 1x Glutamax, 50 U/ml Penicillin-Streptomycin, 50 μM 2-Mercaptoethanol). 14 days after macrophage differentiation, macrophage precursors were harvested using a cell strainer (Stemcell technologies, 27250). Macrophage precursors were added to the NPC culture in 1:1 neural maintenance medium:microglia medium ((DMEM-F12+Glutamax, 100 ng/mL M-CSF (Peprotech, 300-25), 100 ng/mL IL-34 (Peprotech, 200-34), 1x Glutamax, 50 U/ml Penicillin-Streptomycin, 50 μM 2-Mercaptoethanol).

### Replication labelling

Prior to co-seeding of 1321N1 and SH-SY5Y mixed cultures, individual cultures were incubated with either 10 µM EdU (Click-iT® EdU Imaging Kit, Life Technologies, C10340) or 10 µM BrdU (Sigma-Aldrich, B5002) for 24h. This incubation time exceeded the doubling time, allowing incorporation of the nucleotide analog in all cells. This labelling period was followed by a 24h washout period in regular cell culture medium. After washout, the cells were subcultured and plated in coculture. In half of the replicate, SH-SY5Y cells received BrdU while 1321N1 cells received EdU. For the remainder of wells, the pre-label switched cell types.

### Morphological staining

Morphological staining (Cell Painting) was adapted from Bray et al. 2016^19^. After careful titration, all dye concentrations were adjusted and evaluated for compatibility with 4-channel laser and filter combinations available on the confocal microscope (see further). Staining was performed on cell cultures fixed in 4% PFA (roti-histofix 4% paraformaldehyde, Roth, 3105.2) for 15min. Cells were rinsed once with PBS-/- (Life Technologies, 10010015) prior to fixation and 3x 5min post fixation. Staining solutions were prepared fresh before staining in PBS-/- with 0.3% Triton-X-100 (Sigma Aldrich, X100) (**Table 1**). Each staining solution was incubated for 30min on a shaker at RT in the dark. After staining, the cells were washed 1x with PBS -/- and sealed for imaging.

**Table 1.**
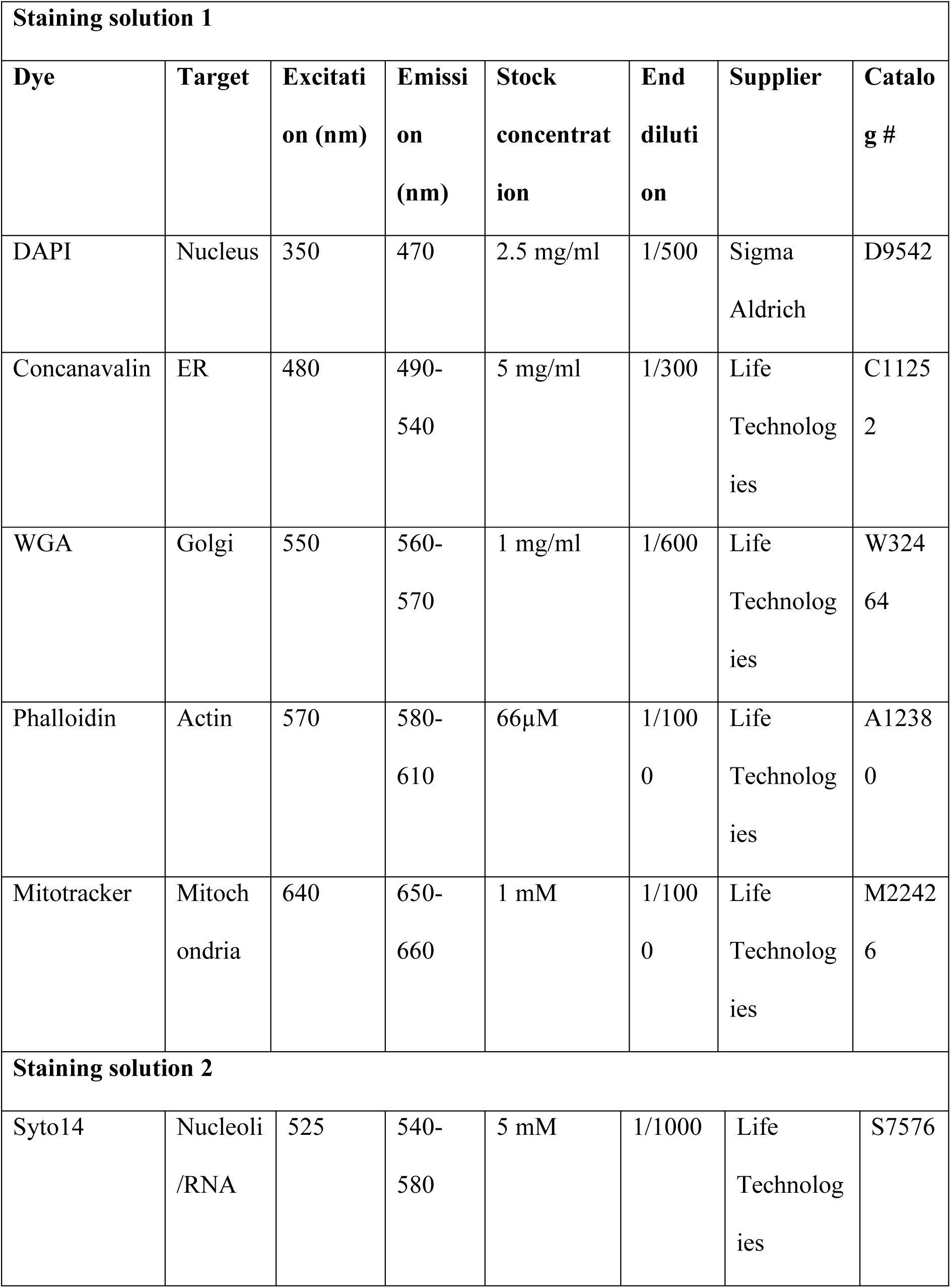
Specifications of morphological staining composition.

### Cyclic staining and immunocytochemistry

Cyclic staining is executed by fluorescence quenching after each sequential imaging round. 1mg/ml in ddH_2_O LiBH_4_ solution^32^ (Acros Organics, 206810050) was prepared fresh before use. 1.5h incubation of quenching solution was performed before each successive staining series. After incubation, the quenching solution was removed by washing 3x 5min in PBS-/-. Successful fluorescence quenching was microscopically verified before proceeding with immunofluorescence staining (IF) (**Table 2, Suppl. Fig. 4**).

**Table 2.** Used antibodies.

| Antibody |  | Host | Supplier | Catalog # |
| --- | --- | --- | --- | --- |
| Primary antibodies | Anti-BrdU | Sheep | Abcam | ab1893 |
|  | Anti-MAP2 | Chicken | Synaptic systems | 188006 |
|  | Anti-TUBB3 | Mouse | BioLegend | 801202 |
|  | Anti-CD44 | Mouse | Millipore | MAB4065 |
|  | Anti-GFAP | Goat | Abcam | ab53554 |
|  | Anti-Iba1 | Rabbit | Wako | 019-19741 |
|  | Anti-Ki67 | Rabbit | Abcam | ab15580 |
| Secondary antibodies | Anti-mouse-Cy3 | Donkey | Jackson ImmunoResearch | 715-165-150 |
|  | Anti-mouse-Cy5 | Donkey | Jackson ImmunoResearch | 715-175-150 |
|  | Anti-sheep-Cy3 | Donkey | Jackson ImmunoResearch | 713-165-147 |
|  | Anti-sheep-Cy5 | Donkey | Jackson ImmunoResearch | 713-175-147 |
|  | Anti-chicken-Cy5 | Donkey | Jackson ImmunoResearch | 703-175-155 |
|  | Anti-goat-Cy3 | Donkey | Jackson ImmunoResearch | 705-165-147 |
|  | Anti-Guinea pig-Cy3 | Donkey | Jackson ImmunoResearch | 706-165-148 |
|  | Anti-Rabbit-Cy5 | Donkey | Jackson ImmunoResearch | 711-175-152 |

Cells are treated with PAV blocking buffer (Thimerosal 0.5% (Fluka, 71230), NaN3 0.1% (Merck, k 6688), Bovine serum albumin (Sigma-Aldrich, A7284), Normal horse serum, PBS-/-) for 8min. The desired primary antibodies (pAB) are diluted in PAV blocking buffer. pAB incubation was performed 12h (overnight) at 4°C after which the cells were washed 1x 5min in PBS-/- followed by incubation in secondary antibody solution (sAB) in PAV + DAPI for nuclear counterstain. sAB staining was performed for 3h at RT while shaking. Prior to imaging, the cells were washed 2x with PBS-/- and stored in PBS-/- + 0,1% NaN_3_. BrdU staining was performed using IF, requiring DNA denaturation before pAB incubation. This was performed by 10 min incubation with 2N HCl at 37°C. HCl was neutralized with 0.1 M sodium borate buffer pH 8.5 for 30min at RT. Cells were washed 3x 5min in PBS-/- before continuing with the general IF protocol. EdU click-it labelling was performed according to the manufacturer’s instructions (Click-iT® EdU Imaging Kit, Life Technologies, C10340) (**Suppl. Fig. 5A**).

### Image acquisition

Images were acquired using a spinning disk confocal microscope (Nikon CSU-W1 SoRa) with a 20x 0.75 NA objective (Plan APO VC 20x DIC N2) and Kinetix sCMOS camera (pinhole 50µm; disk speed 4000 rpm; pinhole aperture 10; bit depth 12-bit, pixel size 0.325µm²). We opted for confocal microscopy instead of widefield to overcome image quality limitations resulting from highly dense cell clusters. 96-well plates were scanned, capturing a minimum of 16 images per well spread in a regular pattern (0,8 mm spacing in x and y) across the surface of the well. If multiple z-slices were acquired (to correct for surface inclinations in the field of view), a maximum projection was performed before further analysis. Morphological images were acquired in all 4 channels. (**Table 3**).

**Table 3.**
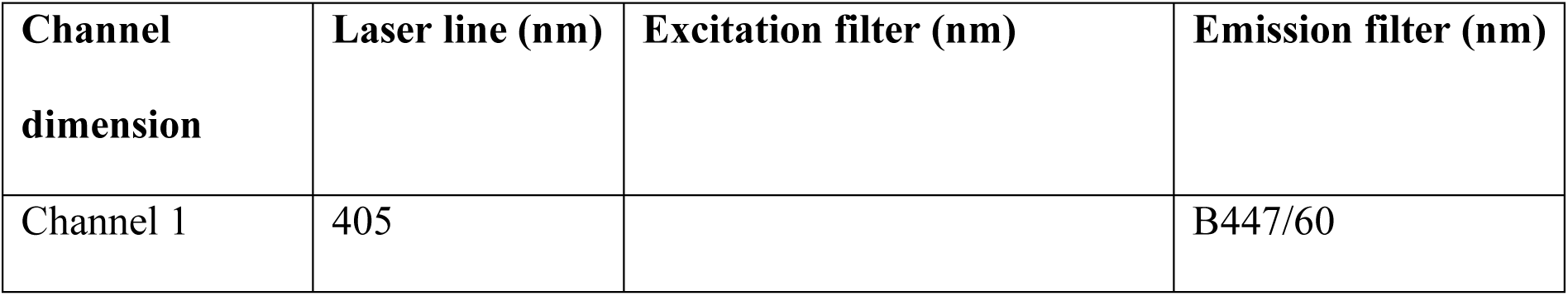
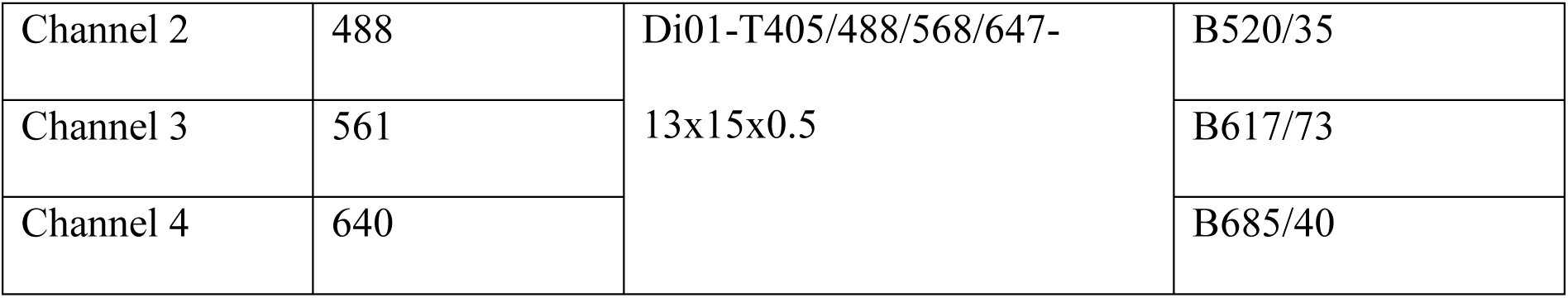
Specifications of the used laser lines, excitation and emission filters.

### Software

Images were captured by the Nikon CSU-W1 confocal system in combination with the NIS elements software (RRID:SCR_014329). Image visualization was later performed using Fiji freeware^51,52^. In-depth image analysis, pre-processing and machine learning for cell classification were performed using Python programming language^53^ in combination with Anaconda^54^ (distribution and package managing software) and Visual Studio Code (code editor). The packages and versions used for data analysis are shown in **Table 4**. **Image pre-processing (Suppl. Fig. 5B).** Note: all image and data analysis scripts are available on GitHub. See further the data availability statement.

**Table 4.**
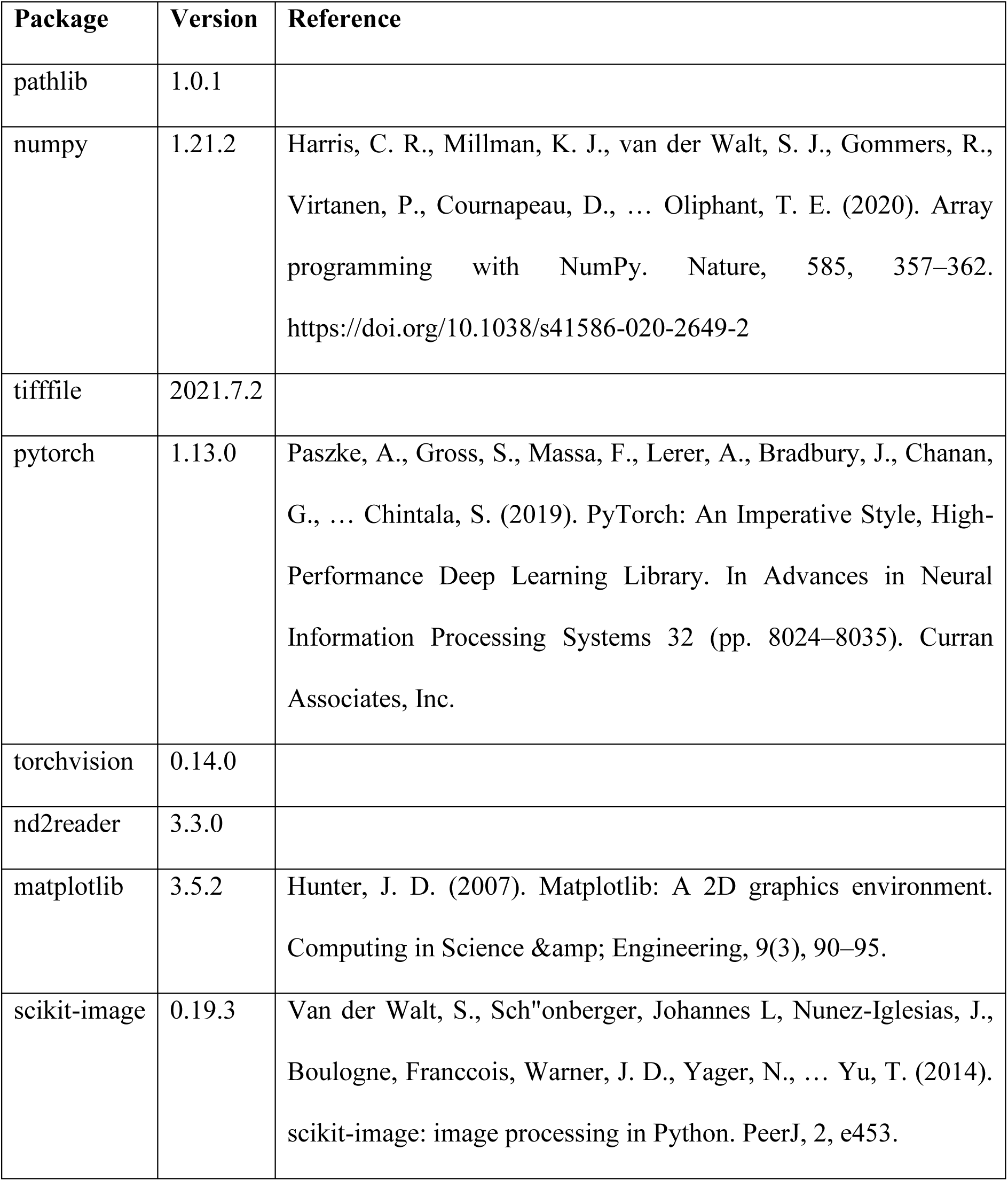
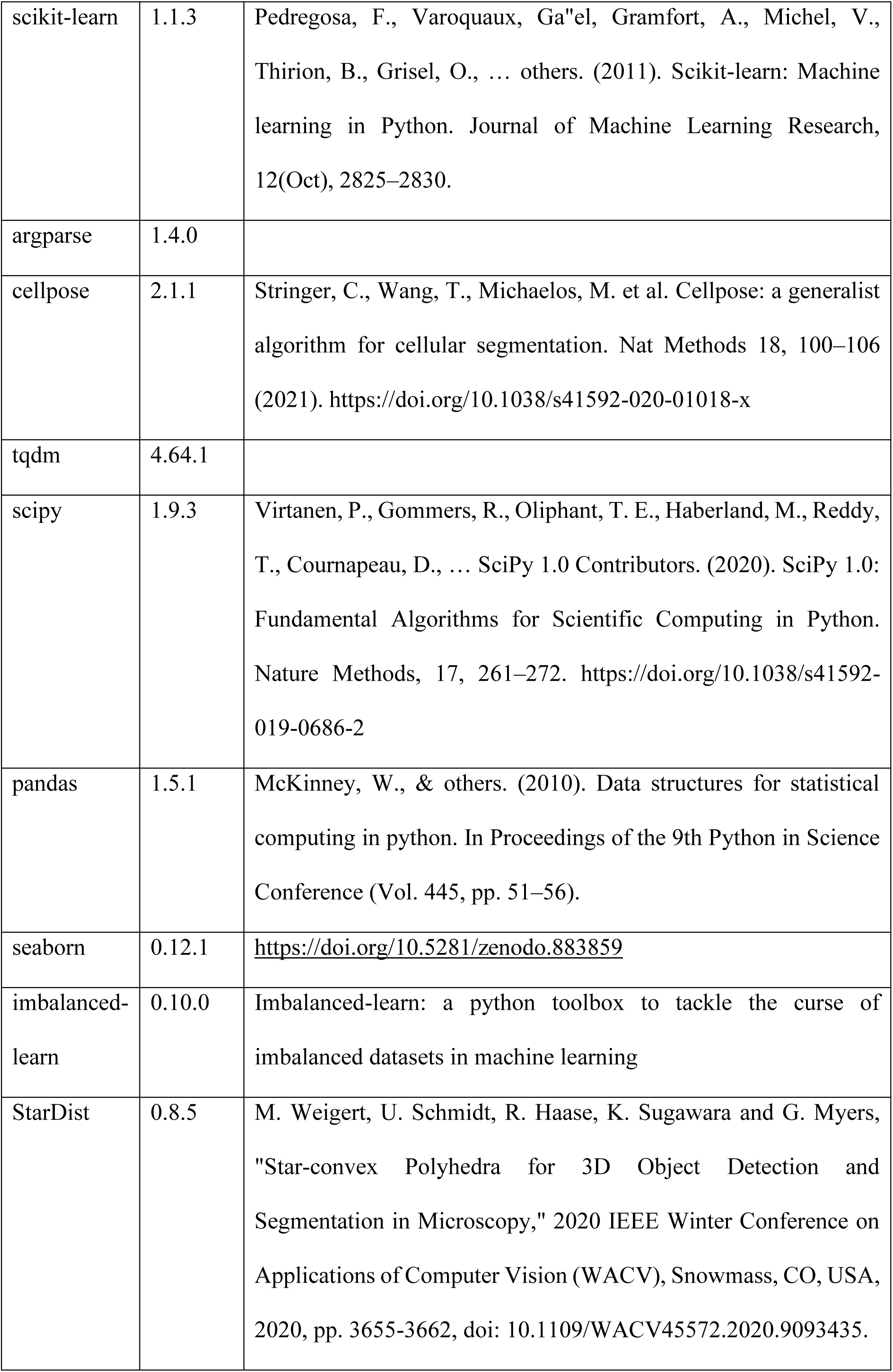
Python packages used for image and data analysis alongside the software version.

#### Cell/nuclear segmentation

Using a tailor-made script implementing the Cellpose^55^ (cell segmentation) or StarDist^56^ (nuclear segmentation) package, images were pre-processed by normalizing the intensity values of each channel between the 1st and 99th quantile. Individual channels or channel combinations for segmentation can be selected depending on the desired outcome mask. Resulting outputs included the segmentation mask alongside the quality control images. For Stardist implementation, hyperparameter probability was set at 0,6 and overlap at 0,3. For Cellpose segmentation, 4 models (cyto2) were averaged to obtain the final mask. Segmentation was performed on the composite of all CP channels. An estimation of the cell’s diameter could be included to optimize cell detection. Cell segmentation was performed using Cellpose and used in all cases where the whole-cell crop was given as input to the CNN or data from the whole cell was used for feature extraction for RF (Fig. 1-4). Nuclear segmentation was performed using Stardist and used for nuclear and nucleocentric crops (Fig. 4-7).

#### Ground truth alignment

Following sequential staining and imaging rounds, multiple images were captured representing the same cell with different markers. Lifting the plate of the microscope stage and imaging in sequential rounds after several days results in small linear translations in the exact location of each image. These linear translations need to be corrected to align morphological with ground truth image data within the same ROI. All images were aligned using Fourier-based image cross correlation on the intensity-normalized multichannel images. The alignment shift between image1 and image2 was determined using scikit-image phase cross correlation. Image2 was then moved according to the predetermined shift to align morphological with ground truth images.

#### Ground truth phenotyping

The true cell phenotype was determined by the fluorescence intensity of the post-hoc immunostaining with class-specific markers or pre-labelling with Edu/Brdu. In this latter case, the base analogues were incorporated into each cell line prior to mixing them, i.e. when they were still growing in monoculture so they could be labelled and identified after co-seeding and morphological profiling. Each ground truth image was imported alongside the corresponding cell mask for that image. For each cell label, the fluorescence intensity was determined and tabulated. The threshold was set manually based on the fluorescence intensity of the monoculture controls. Ground truth labels were assigned to each region of interest (ROI).

#### Feature extraction

Handcrafted features were extracted using the scikit-image package (regionprops and GLCM functions). The definition of each feature extracted from the image is listed in **table 5**. All features were extracted for each channel in the cell painting image and for every region (cell, cytoplasm and nucleus) within the ROI.

**Table 5.** Definition of handcrafted features (according to the scikit-image documentation). All of these features were extracted for each fluorescent channel (1-4) and region (nucleus, cytoplasm and whole-cell) in the cell painting images.

| Feature name | Description | Category |
| --- | --- | --- |
| Intensity max | Maximal pixel intensity inside the ROI | Intensity |
| Intensity mean | Mean pixel intensity inside the ROI | Intensity |
| Intensity min | Minimal pixel intensity inside the ROI | Intensity |
| Intensity Std | Standard deviation of the pixel intensities inside the ROI | Intensity |
| Area | Area of the ROI | Shape |
| Area convex | The area of the convex polygon that encloses the ROI | Shape |
| Area filled | Area of the ROI including all holes inside the ROI | Shape |
| Axis major length | The length of the major axis of the ellipse that has the same normalized second central moments as the region. | Shape |
| Axis minor length | The length of the minor axis of the ellipse that has the same normalized second central moments as the region. | Shape |
| Centroid | Coordinates of the center of the ROI | Shape |
| Eccentricity | Eccentricity of the ellipse that has the same second-moments as the ROI. | Shape |
| Equivalent diameter area | The diameter of a circle with the same area as the ROI. | Shape |
| Extent | The ratio of the Area of the ROI to the Area of the bounding box around the ROI. | Shape |
| Feret diameter max | The longest distance between points around a ROIs' convex polygon. | Shape |
| Orientation | The angle between the X-axis and the major axis. | Shape |
| Perimeter | Length of the contour of the ROI | Shape |
| Perimeter crofton | Perimeter of the ROI approximated by the Crofton formula in 4 directions. | Shape |
| Solidity | The ratio of the number of pixels inside the ROI to the number of pixels of the convex polygon. | Shape |
| Contrast | The difference between the maximum intensity and minimum intensity pixel inside the ROI. | Texture |
| Dissimilarity | The average difference in pixel intensity between neighbouring pixels. | Texture |
| Homogeneity | Value for similarity of pixels in the ROI. | Texture |
| Energy | Value for the local change of pixel intensities in the ROI. | Texture |
| Correlation | Value for the linear dependency of pixels in the ROI. | Texture |
| ASM (angular second moment) | Value for the uniformity of pixel values in a ROI. | Texture |

#### Data filtering

Over the entire pipeline, ROIs could be discarded based on 3 conditions: (1) Cell detachment. Due to the harsh sample preparation and repeated washing steps, cells could detach from the imaging substrate thus resulting in lack of ground truth information for those cells. As a result, all ROIs for which there was no DAPI signal detected in the ground truth image, were removed from the dataset as incomplete. (2) Cells for which the ground truth fluorescence intensity was ambiguous. Ground truth labels were determined based on the presence of specific IF staining in either of the phenotype classes. If no class-specific IF staining was detected by thresholding, no true label could be assigned. These ROIs were therefore discarded due to uncertainty. (3) Likelihood of faulty ROI detection. The DAPI signal was used to discard ROIs that do not represent (complete) cells by thresholding on minimal DAPI intensity (mean nuclear DAPI intensity > 500) and minimal nuclear size (nuclear area > 160).

### ROI classification

#### Train-validation-test split

Model training requires splitting the available data in 3 subsets: training (60%), validation (10%), and testing (30%). The training dataset was used to train the machine learning models (either RF or CNN). The validation dataset, composing of 10% of the total data, was used to determine the hyperparameters and intermediate testing. The remaining 30% of the dataset was kept apart and used to test the final model when training is completed. For both RF and CNN, the testing dataset was never shown to the model during the training phase, but only used after training to determine the accuracy of predictions to the ground truth. The number of instances for each class was equalized by sampling equal number of instances from each predicted class. To account for variation between technical replicates, the train-validation-test split was stratified per well. As a result, no datapoints arising from the same well could appear simultaneously in the training, validation and testing subset. This data stratification was repeated 3 times with different random seeds (see Methods – Reproducibility).

#### Random Forest

For each ROI, a set of manually defined parameters was extracted corresponding to cell shape (area, convex area, filled area, length minor axis, length major axis, centroid, eccentricity, equivalent diameter area, ferret diameter max, orientation, perimeter, solidity), texture (GLCM: contrast, dissimilarity, homogeneity, energy, correlation, ASM) and intensity (maximal intensity, minimal intensity, mean intensity, standard deviation intensity). This was done for 3 regions (nucleus, cytoplasm and complete cell), and for all channel dimensions. Redundant parameters could be removed if above a 0.95 correlation threshold. All parameters were standardized per ROI, grouped per replicate. The number of trees within the forest was varied between 10 and 150, reaching maximum accuracy at around 30 trees.

#### Uniform Manifold Approximation and Projection

Dimensionality reduction using UMAP was performed using either the same feature matrix as used for RF prediction or the feature embeddings from the trained CNN classification network. Hyperparameters were set at the default settings.

#### Convolutional neural network

A ResNet50 model was trained for image classification. In contrast to classical machine learning techniques, no handcrafted features were extracted. Crops were defined based on the segmentation mask for each ROI. The bounding box was cropped out of the original image with a fixed patch size (60µm for whole cells, 18µm for nucleus and nucleocentric crops) surrounding the centroid of the segmentation mask. For the whole cell and nuclear crops, all pixels outside of the segmentation mask were set to zero. This was not the case for the nucleocentric crops. Each ROI was cropped out of the original morphological image and associated with metadata corresponding to its ground truth label. Images alongside their labels were fed to the network. Tensors were normalized per channel. Models are trained on a minimum of 5000 training inputs of each class for 50 epochs (training iterations). Each batch consisted of 100 samples. The training input was augmented by linear transformations (horizontal and vertical flip, random rotation). Each epoch, the current model was tested against a validation dataset (**Suppl. Fig. 5C**). The performance of the model on this validation subset determined whether the model was stored as new best (if the new accuracy exceeded the accuracy of the previous best model) or discarded. The learning rate at the start was set at 0,0001 and automatically reduced with a factor of 0,1 during training when no improvement was seen after 10 epochs. After 50 epochs, the best resulting model was tested on a separate test dataset to determine the accuracy on previously unseen data.

### GradCAM

GradCAM analysis was used to visualize the regions used by the CNN for classification. This map is specific to each cell. Images are selected randomly out the full dataset for visualization. To avoid cherry-picking, a set of GradCam maps is reported alongside the random seed used for image selection.

### Reproducibility

Each model training was performed 3 independent times (model initializations, repetitions are indicated with ‘N’ on the figures). This was repeated for 3 different random seeds. Each model received input data arising from a minimum of 16 images per well, at least 15 technical replicates (wells). The optimization experiments (figures 1-4) were performed with cell lines with limited variability. These models were trained on 3 independent experiments where ground truth pre-labelling (Edu/BrdU) was performed at least once on either of the cell lines in coculture. For iPSC-derived cultures, as variability is inherent to these differentiations, 3 biological replicates (independent differentiations) were pooled for model training.

### Statistics

All statistical comparisons were made nonparametric using Mann-Whitney U (for two independent sample comparison) or Kruskal-Wallis (for multiple sample comparison) with pairwise tests using Tukey’s honestly significant difference test. We opted for nonparametric testing because the number of models in each group to be compared was < 15. Significance levels are indicated on the figures using ns. (no statistical significance, p-value above 0,05), * (p-value between 0,05 and 5e-4), ** (p-value between 5e-4 and 5e-6) and *** (p-value smaller than 5e-6).

## Acknowledgements

We thank Marlies Verschuuren and Hugo Steenberghen for their assistance and knowledge regarding the cell lines. By extension, we would like to thank all current and former members of the De Vos lab.

## Data availability statement

The authors report that the results of this study are available within the manuscript and supplementary materials. All image analysis scripts are open-source available on GitHub (https://github.com/DeVosLab/Nucleocentric-Profiling)

## Abbreviations

BrdU: Bromodeoxyuridine
CNN: convolutional neural network
CP: cell painting
DAPI: 4’,6- diamidino-2-fenylindool
DIV: days in vitro
EdU: 5-ethynyl-2’-deoxyuridine
ER: endoplasmic reticulum
Grad-CAM: Gradient-weighted Class Activation Mapping
IF: immunofluorescence
iPSC: induced pluripotent stem cell
NPC: neural progenitor cell
RF: random forest
ROI: region of interest
TUBB3: beta-III-tubulin
UMAP: Uniform Manifold Approximation and Projection

## Author contributions

The experiments were conceptualized by SDB, TVDL, JVDD, PP and WDV. Experiments were executed by SDB and JVDD. TVDL and SDB optimized the data analysis scripts. SDB analysed the imaging data and performed the data analysis. SDB and WDV prepared the original draft. PP and WDV supervised the work. All authors took part in reviewing and editing the manuscript.

---

**Supplementary figure 1.**
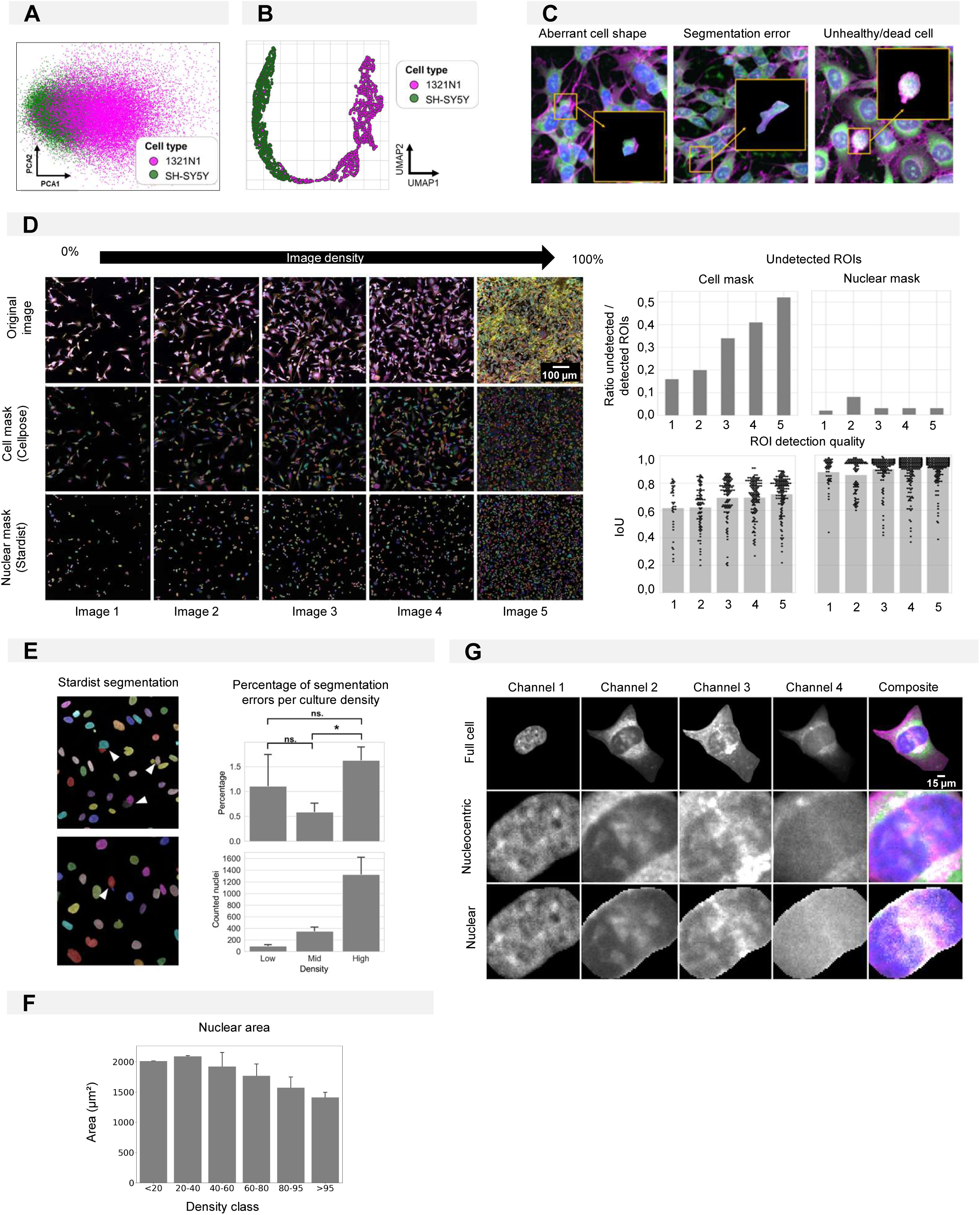
(A) PCA dimensionality reduction on handcrafted features extracted from monocultures of 1321N1 and SH-SY5Y cells. Each point represents an individual cell. (B) UMAP of feature embeddings of the CNN trained to classify 1321N1 and SH-SY5Y monocultures. Each point represents an individual cell. (C) Examples of misclassified ROIs. (D) Left: Representative images of 1321N1 cells with increasing density alongside their cell and nuclear mask produced using resp. Cellpose and Stardist. Images are numbered from 1-5 with increasing density. Upper right: The number of ROIs detected in comparison to the ground truth (manual segmentation). A ROI was considered undetected when the intersection over union (IoU) was below 0,15. Each bar refers to the image number on the left. The IoU quantifies the overlap between ground truth (manually segmented ROI) and the ROI detected by the segmentation algorithm. It is defined as the area of the overlapping region over the total area. IoU for increasing cell density for cell and nuclear masks is given in the bottom right. Each point represents an individual ROI. Each bar refers to the image number on the left. (E) Examples of segmentation mistakes made by the Stardist segmentation algorithm for nuclear segmentation for different culture densities. (F) Nuclear area in function of density. (G) Definition of cell regions given as training input for nuclear and nucleocentric model training.

**Supplementary figure 2.**
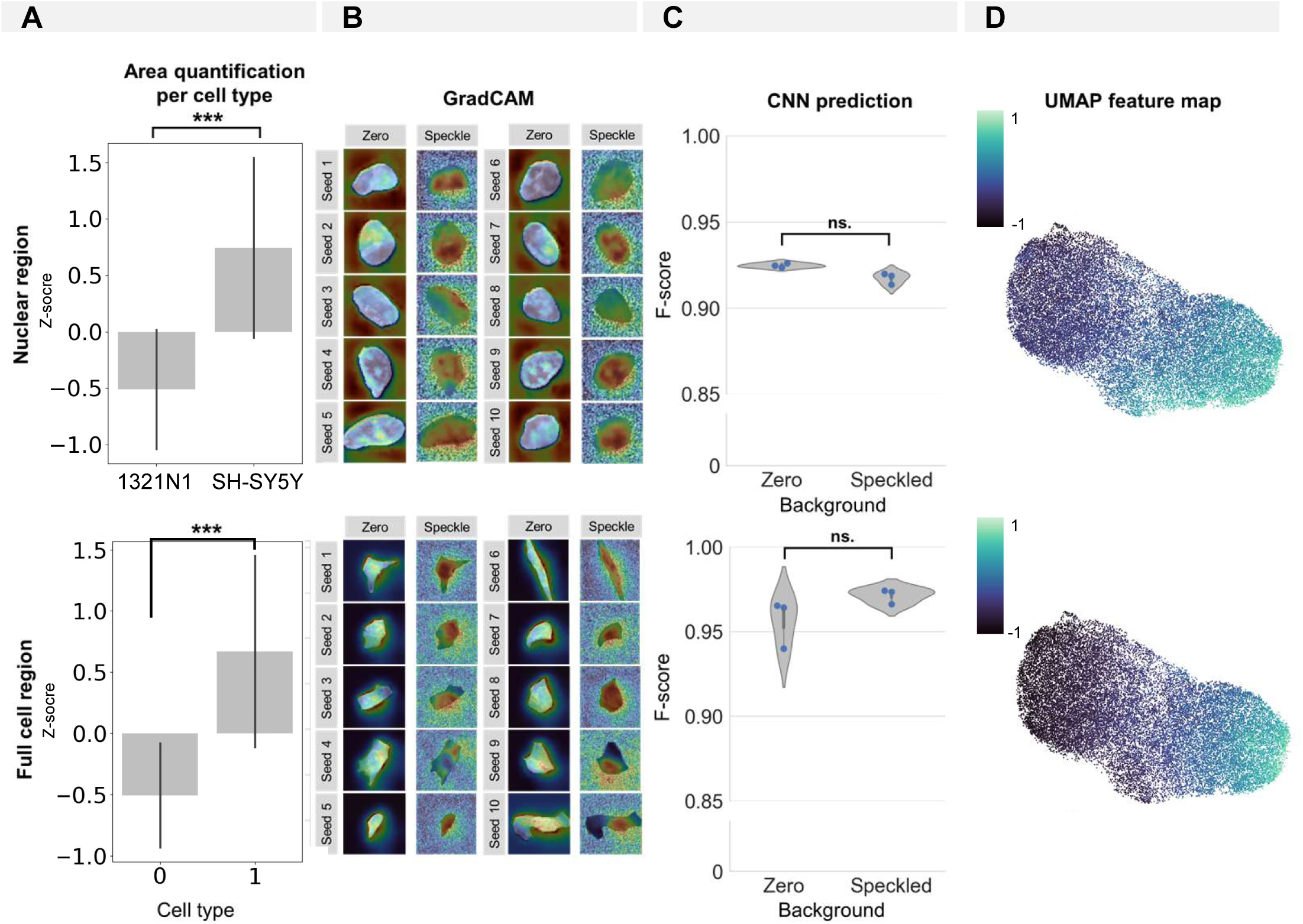
Influence of the nuclear/cellular size on CNN prediction and association with the input background (space artificially set to zero outside of the segmentation mask). (A) Quantification of average nuclear/cellular size per cell type. (B) GradCAM images for 10 random seeds for crops and CNN models trained with background either set to zero or ‘random speckle’. (C) CNN prediction results for models trained on crops with background either set to zero or ‘random speckle’. Each dot in the violinplots represents the F-score of one classifier (model initialization, N = 3). (D) Feature map (see UMAP in figure 1 B and C) of 1321N1 and SH-SY5Y cells showing the contribution of nuclear/cellular area to the cell type cluster separation. Each point represents an individual cell.

**Supplementary figure 3.**
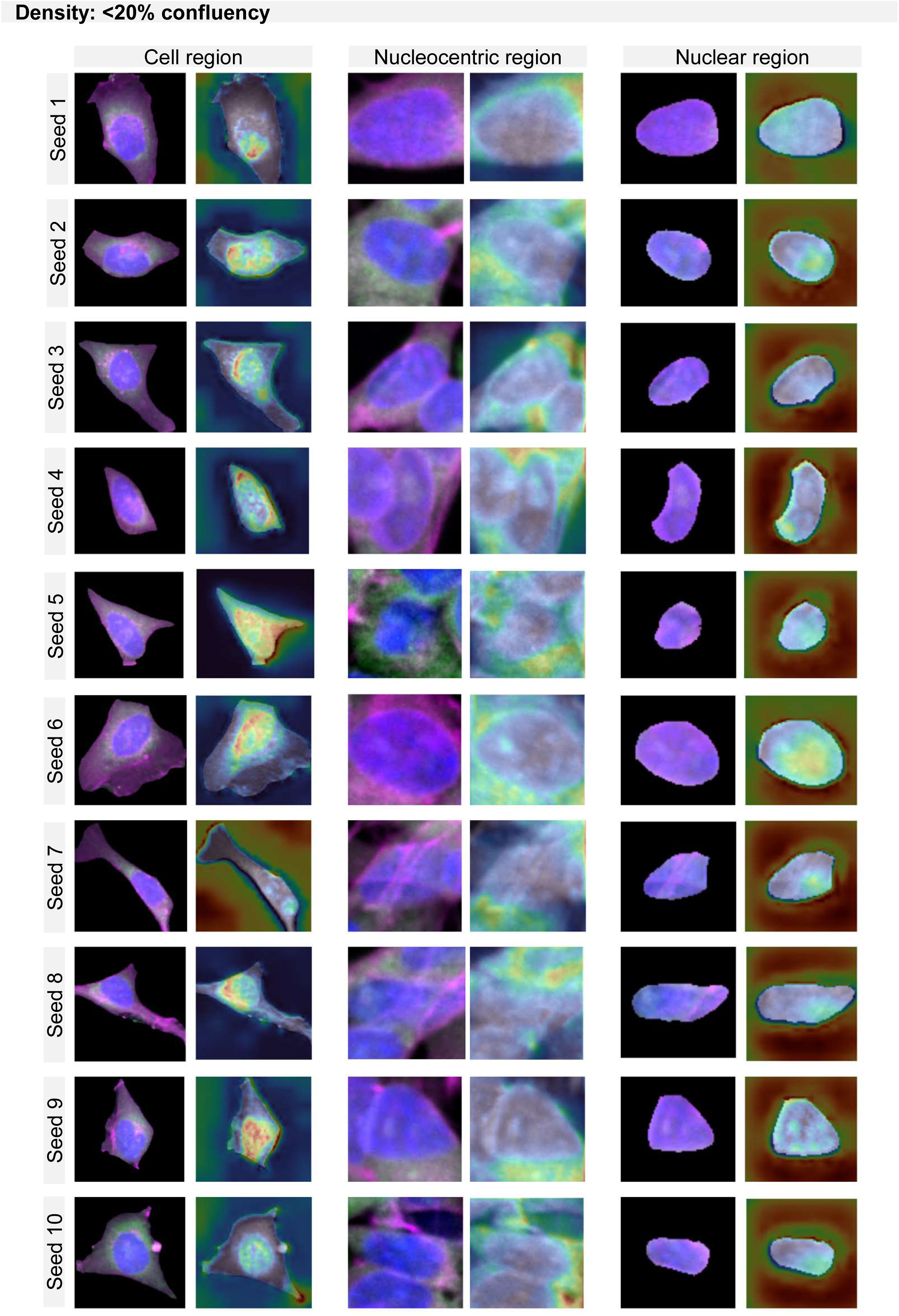
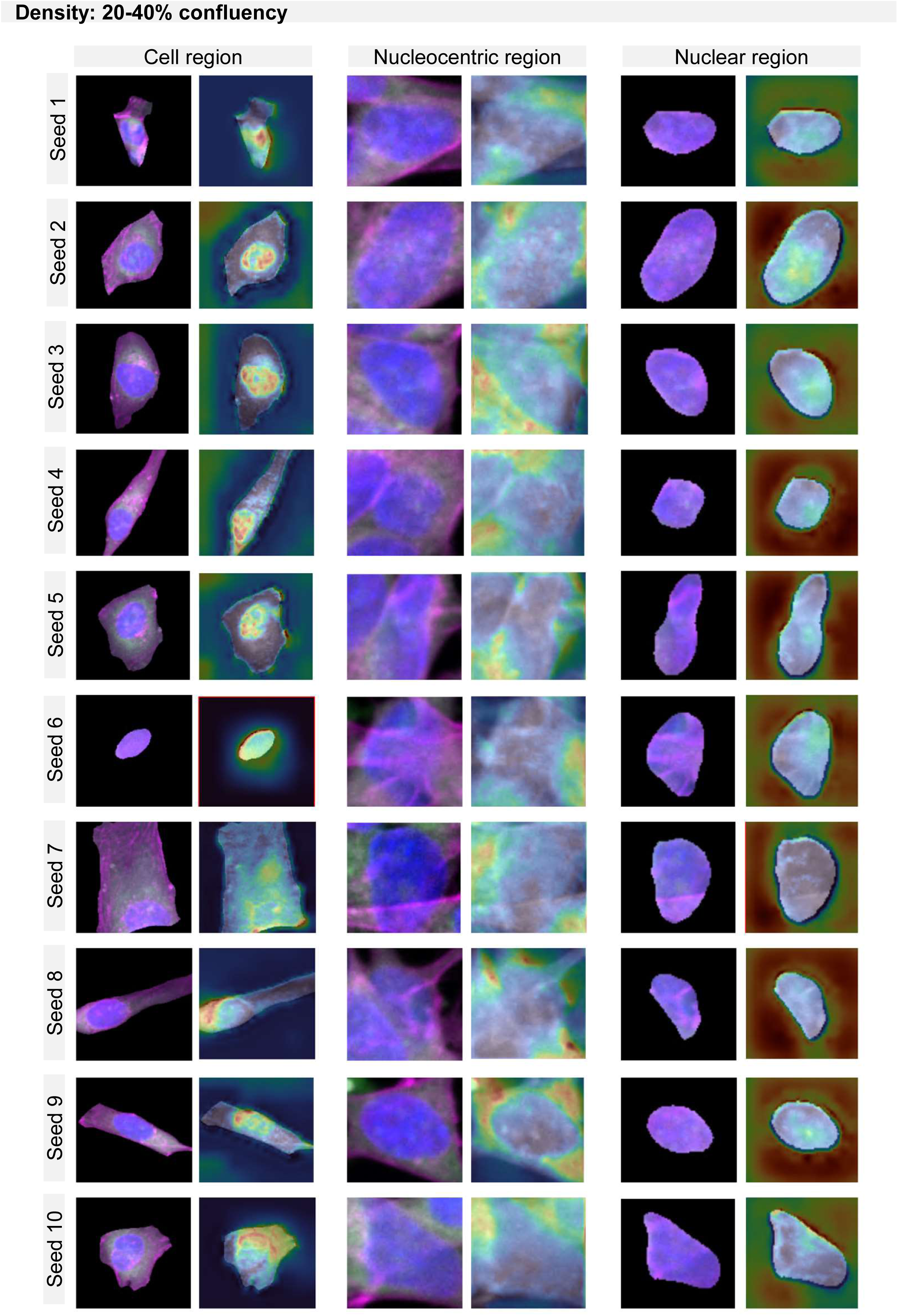
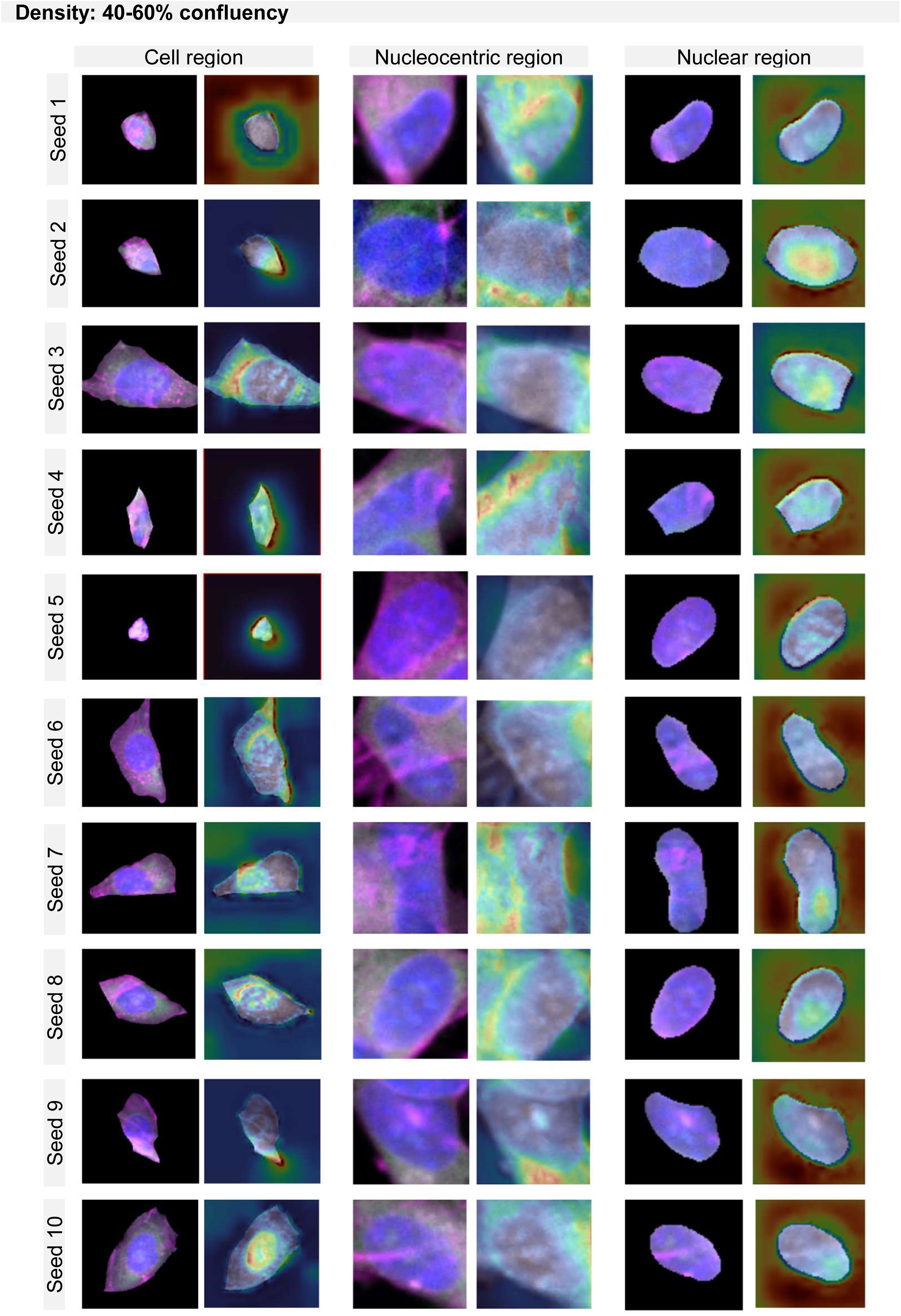
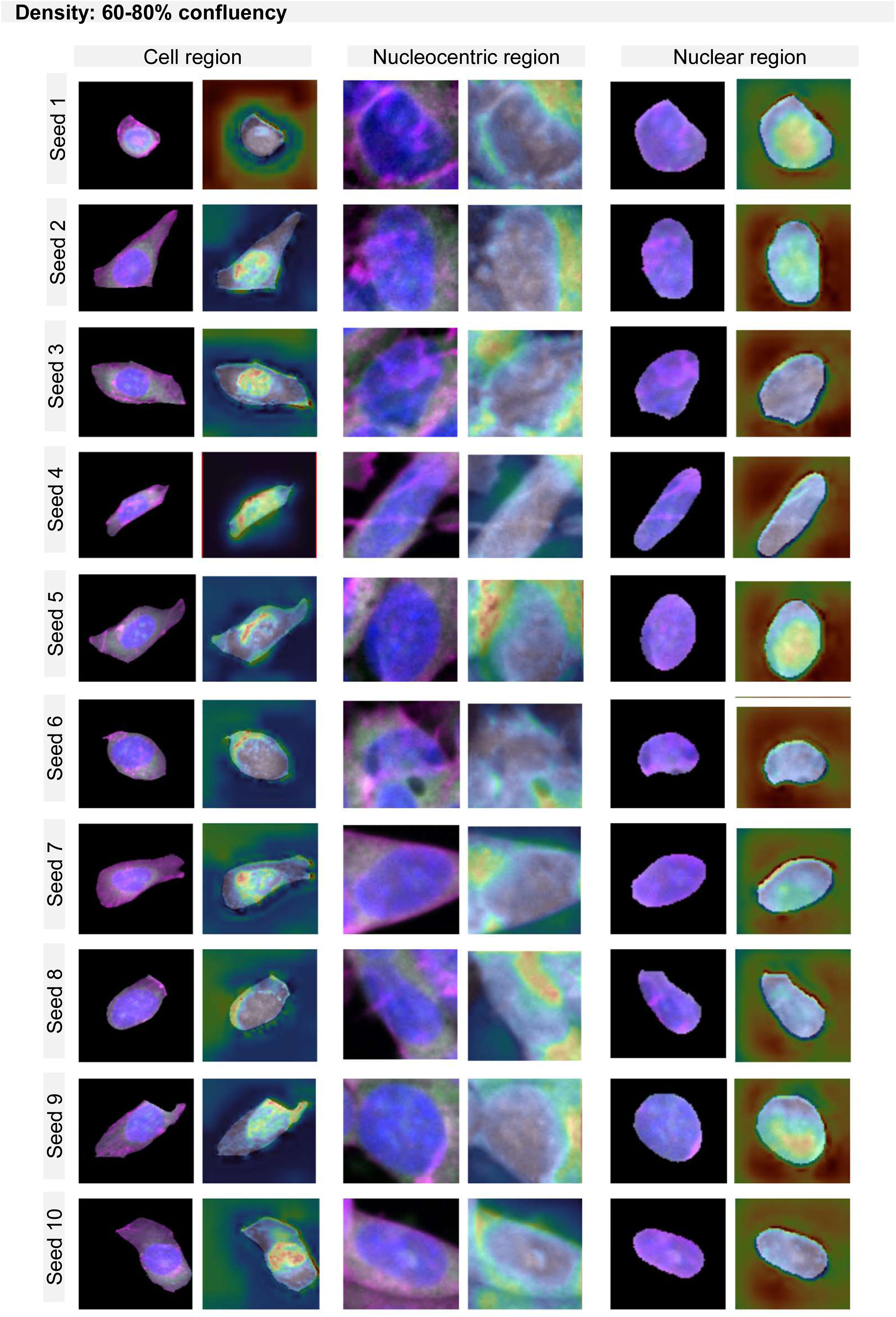
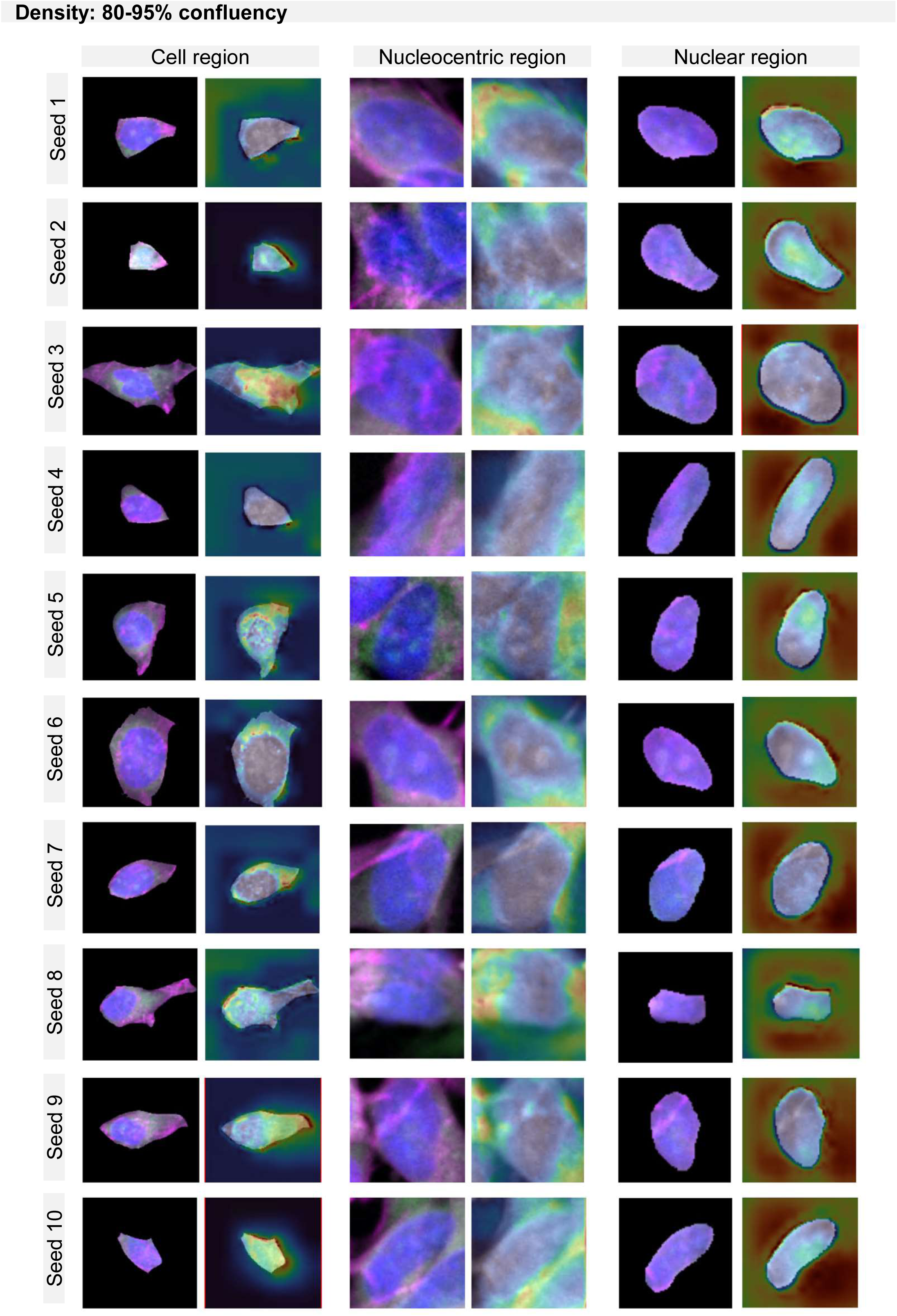
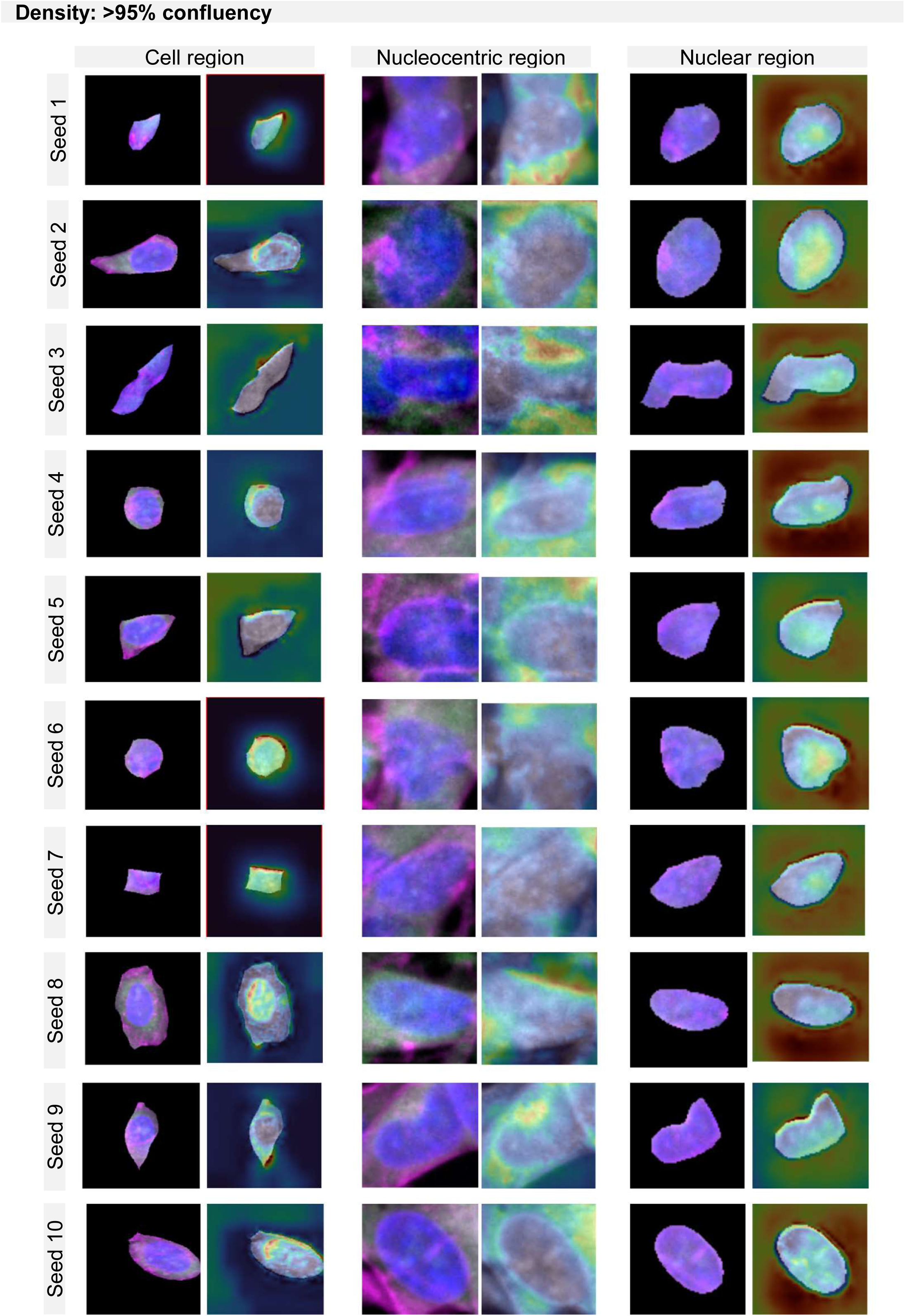
GradCAM maps per region and density.

**Supplementary figure 4.**
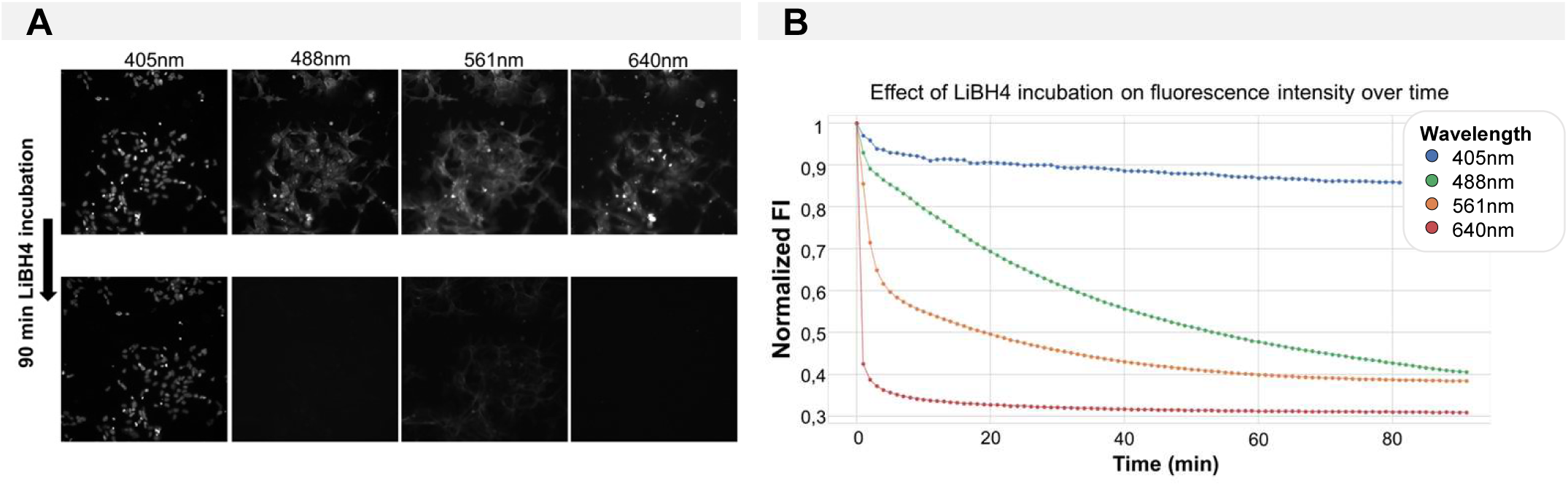
Fluorescence quenching over time using LiBH4. (A) Images before and after quenching for all 4 fluorescence channels. (B) Time curve of normalized image-level fluorescence intensity during incubation with 1mg/ml LiBH4.

**Supplementary figure 5.**
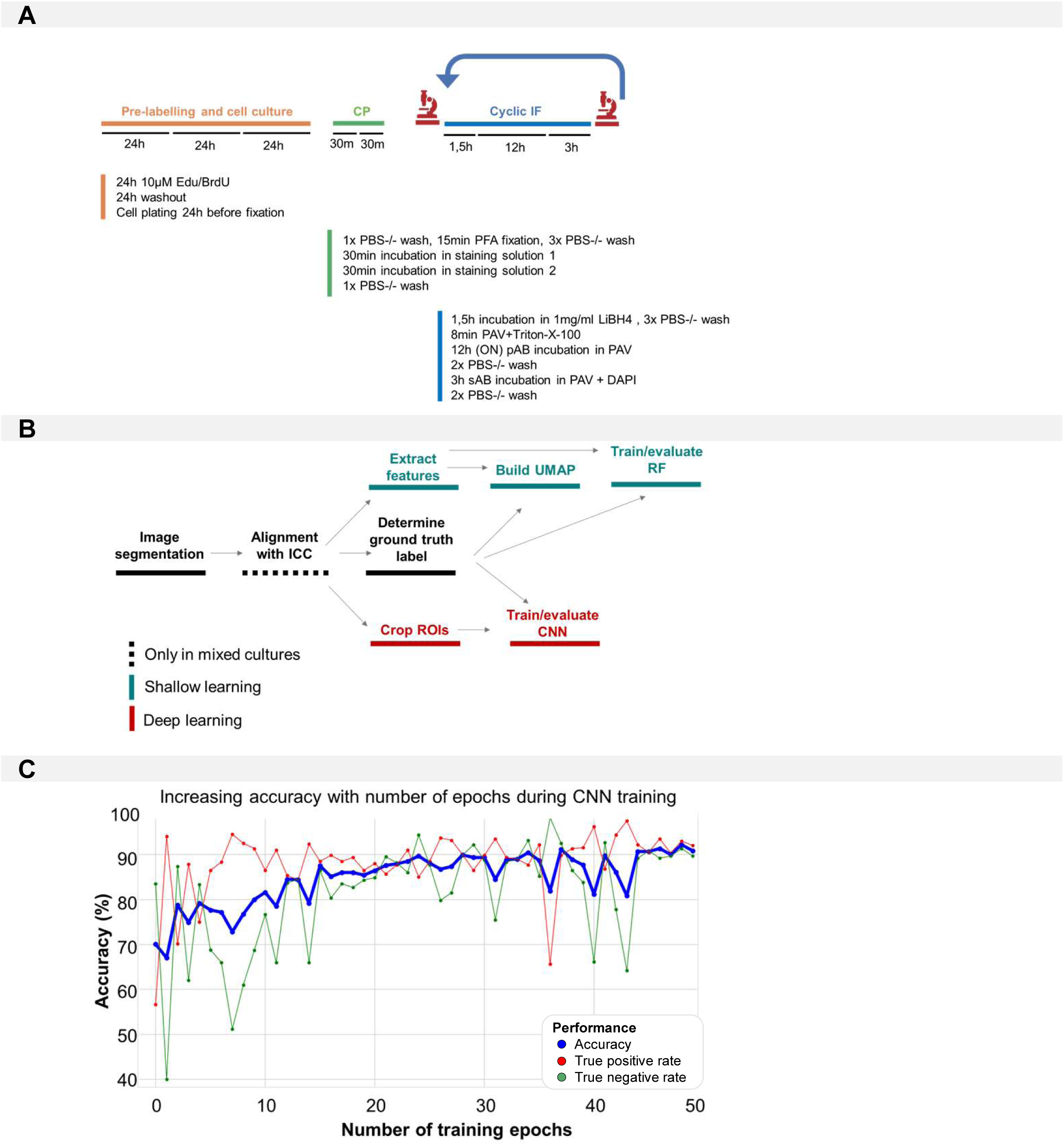
Methods. (A) Pipeline used for morphological profiling in mixed cultures. (B) Overview of the steps within the image analysis pipeline. (C) Evaluation of accuracy, true negative and true positive rate during CNN training (1321N1 vs. SH-SY5Y in monocultures) across all 50 epochs.

